# Slit2 is necessary for optic axon organization in the zebrafish ventral midline

**DOI:** 10.1101/2020.09.25.314062

**Authors:** Camila Davison, Flavio R. Zolessi

## Abstract

Slit-Robo signaling has been implicated in regulating several steps of retinal ganglion cell axon guidance, with a central role assigned to Slit2. We report here the phenotypical characterization of a CRISPR-Cas9-generated zebrafish null mutant for this gene, along with a detailed analysis of its expression pattern by WM-FISH. All evident defects in the optic axons in *slit2-/-* mutants were detected outside the retina, coincident with the major sites of expression at the ventral forebrain, around the developing optic nerve and anterior to the optic chiasm/proximal tract. Anterograde axon tracing experiments in zygotic and maternal-zygotic mutants, as well as morphants, showed the occurrence of axon sorting defects, which appeared mild at the optic nerve level, but more severe in the optic chiasm and the proximal tract. A remarkable sorting defect was the usual splitting of one of the optic nerves in two branches that surrounded the contralateral nerve at the chiasm. Although all axons eventually crossed the midline, the retinotopic order appeared lost at the proximal optic tract, to eventually correct distally. Time-lapse analysis demonstrated the sporadic occurrence of axon misrouting at the chiasm level, which could be responsible for the sorting errors. Our results support previous evidence of a channeling role for Slit molecules in retinal ganglion cell axons at the optic nerve, in addition to a function in the segregation of axons coming from each nerve and from different retinal regions at the medio-ventral area of the forebrain.

## 1 INTRODUCTION

The neural retina of vertebrates develops as an evaginated extension of the early forebrain, to which it remains connected through the optic stalk. Hence, the axons of its projection neurons, the retinal ganglion cells (RGCs), must undergo a long and highly regulated process of guided growth to eventually reach their synaptic targets in the mesencephalic roof. RGCs differentiate in the basal-most cell layer of the retina, and grow an axon from their basal poles, to later develop a dendritic tree from the apical region (Zolessi et al., 2006). Along their journey, RGC growth cones encounter several different signaling molecules that guide their path (Herrera et al., 2019). The first portion is particularly challenging, as it includes intraretinal guidance to find the optic nerve exit, the turning and fasciculation to form the optic nerve, the exit from the eye, growth along the optic stalk, and crossing (or not, depending on species) at the optic chiasm. A family of signaling molecules, the Slit proteins, and their Robo receptors, have been found to be relevant in these events, largely acting as repulsive factors, although their exact mechanisms of action are still not completely understood (Herrera et al., 2019).

Slit molecules constitute a family of large, multi-modular, secreted glycoproteins of ≈200 kDa, with various forms present across animals with bilateral symmetry (Brose and Tessier-Lavigne, 2000). Invertebrates like *D. melanogaster* have only one form of this protein (named Slit), while there are three in most vertebrates (Slit1-3) and four in the zebrafish (Slit1a, 1b, 2 and 3) (Hutson et al., 2003; Yeo et al., 2001). Although Slit ligands are secreted, their diffusion is limited due to their strong association with extracellular matrix components such as collagen IV and dystroglycan (Wright et al., 2012; Xiao et al., 2011). Slit main receptors are the Robo family of proteins, to which they bind through the second LRR domain (Howitt et al., 2004). There are four genes coding for these receptors in vertebrates, including the zebrafish (Robo1-4) (Lee et al., 2001; Park et al., 2003).

Slit2 was shown to be important for intraretinal axon pathfinding in mice (Thompson et al., 2006), and to act together with Slit1 to channel RGC axons to their correct location at the optic chiasm (Plump et al., 2002). In the zebrafish, RGCs strongly express *robo2* around the time they extend their axons and its null mutant *astray*, presents severe defects in axon guidance outside the retina (Fricke et al., 2001). Slits mRNAs have been reported to be differentially expressed along the optic pathway in the zebrafish embryo: *slit1a, slit1b* and *slit2* appear in the retina, with the latter apparently being expressed later and just in some cells of the inner nuclear layer, while *slit1a*, *slit2* and *slit3* are expressed around the optic nerve and optic chiasm area, with different distributions and relative abundance (Barresi et al., 2005; Chalasani et al., 2007; Hutson and Chien, 2002; Wong et al., 2012; Yeo et al., 2001). In the optic nerve/chiasm area in particular, *slit2* expression was found within the optic stalk and surrounding the optic nerve, as well as in the rostral margin of the optic recess, which is located midway between the anterior commissure and post-optic commissure, and in close proximity to the optic chiasm (Barresi et al., 2005; Chalasani et al., 2007; Hutson and Chien, 2002), all indicating a potentially important role for this secreted factor in RGC axon guidance around this area.

Here, we describe a new *slit2* mutant in the zebrafish (NM_131753.1:g.30_39del), generated through CRISPR-Cas9, and present an initial characterization of its phenotype regarding RGCs and their axons. We found that Slit2 is important for RGC axon bundling at the optic nerve and sorting at the optic chiasm and proximal tract, while apparently not being necessary at the distal optic tract and optic tectum.

## 2 MATERIALS AND METHODS

### 2.1 Fish breeding and care

Zebrafish were maintained and bred in a stand-alone system (Tecniplast, Buguggiate, Italy), with controlled temperature (28 °C), conductivity (500 μS/cm^2^) and pH (7.5), under live and pellet dietary regime. Embryos were raised at temperatures ranging from 28.5 to 32 °C and staged in hours post-fertilization (hpf) according to Kimmel and collaborators (Kimmel et al., 1995). Fish lines used: wild-type (Tab5), Tg(*atoh7*:gap43-EGFP)^cu1^ (Zolessi et al., 2006), SoFa1 (*atoh7*:gap-RFP/ptf1a:cytGFP/*crx*:gap-CFP; Almeida et al., 2014), and the mutant line generated here (see below) NM_131753.1:g.30_39del (hereafter denoted as *slit2-/-*). All the manipulations were carried out following the approved local regulations (CEUA-Institut Pasteur de Montevideo, and CNEA).

### 2.2 Generation of the slit2-/- line

We designed four single-guide RNAs (sgRNA) against the *slit2* gene using the CRISPRscan tool (Moreno-Mateos et al., 2015; Supplementary Table 1). We then tested them individually by injecting each one together with mRNA for zfCas9 flanked by two nuclear localization signal sequences (nCas9n; Jao et al., 2013)). One of them, complementary to a sequence in the first exon of the coding sequence (sgRNA *slit2* 21; Supplementary Table 1), presented no toxic effects and was highly efficient, as evidenced by agarose gel electrophoresis. We injected one-cell stage Tab5 embryos with this sgRNA, together with Cas9 mRNA, and raised them to adulthood. A chosen female was subjected to outcross with a wild-type male, and the progeny was genotyped at 48 hpf. After gel electrophoresis and sequencing, we found a set of four different mutations (Supplementary Table 2). The embryos obtained from the outcross were raised to adulthood, and were then selected for the desired 10 base pair deletion through fin clipping. This gave rise to the heterozygous line, which was crossed to obtain the *slit2-/-* mutant NM_131753.1:g.30_39del.

### 2.3 Morpholino treatment

The morpholino oligomers (MOs) used in this study were obtained from Gene Tools (Philomath, USA). The *slit2* MO has been previously used to target zebrafish *slit2* translational initiation: *slit2*-ATG (Supplementary Table 1; Barresi et al., 2005). For *ptf1a* knock-down, a combination of two MOs was used: a translation blocking *ptf1a* MO1 and a splice-blocking *ptf1a* MO2 (Supplementary Table 1; Almeida et al., 2014). 1.5 ng of the *slit2* MO or, alternatively, a mix of 2 ng each of the *ptf1a* MO1+MO2 were injected in the yolk of 1–4 cell-stage embryos, at a maximum volume of 2 nL. As control, we used matching doses of a standard control MO, and we included in all cases an anti-p53 MO to prevent non-specific cell death (Supplementary Table 1), both obtained from Gene Tools (Philomath, USA) and used as previously described (Lepanto et al., 2016).

### 2.4 RT-PCR

Total RNA was extracted from oocytes (n=50), 4 hpf (n=40) or 48 hpf (n=25) embryos using the Aurum Total RNA Mini Kit (Bio-Rad). cDNA was then prepared with the RevertAid reverse transcriptase (Thermo Fisher), using both oligo dT and random hexamer primers. The cDNA was then used as PCR template to amplify either a 182 bp fragment of the *elongation factor 1a* (*ef1a*), used as a control, or a 255 bp region of *slit2*. Primer sequences are shown in Supplementary Table 1.

### 2.5 Whole-mount immunofluorescence

Embryos were grown in 0.003 % phenylthiourea (Sigma, St. Louis, USA) from 10 hpf onwards to delay pigmentation, and fixed overnight at 4 °C, by immersion in 4 % paraformaldehyde in phosphate buffer saline, pH 7.4 (PFA-PBS). For whole-mount immunostaining all subsequent washes were performed in PBS containing 1 % Triton X-100. Further permeability was achieved by incubating the embryos in 0.25 % trypsin-EDTA for 10–15 min at 0 °C. Blocking was for 30 min in 0.1 % bovine serum albumin (BSA), 1 % Triton X-100 in PBS. The primary antibodies, diluted in the blocking solution, were used as follows: zn8 (ZIRC, Oregon, USA), recognizing the adhesion molecule neurolin/DM-grasp, 1/100; anti-acetylated α-tubulin (Sigma, St. Louis, USA), 1/1000; anti-neurofilament-associated antigen (3A10, DSHB), 1/100. The secondary antibody used was anti-mouse IgG-Alexa 488 (Life Technologies, Carlsbad, USA), 1/1000. When necessary, TRITC-conjugated phalloidin (Sigma, St. Louis, USA) was mixed with the secondary antibody. Nuclei were fluorescently counterstained with methyl green (Prieto et al., 2014). All antibody incubations were performed overnight at 4 °C. Embryos were mounted in 1.5 % agarose-50 % glycerol in 20 mM Tris buffer (pH 8.0) and stored at 4 °C or −20 °C. Observation of whole embryos was performed using a Zeiss LSM 880 laser confocal microscope, with a 25X 0.8 NA glycerol immersion objective.

### 2.6 Cryosections

Five-day-old embryos were fixed as described above, washed in PBS and cryoprotected by a 30 min incubation in 5 % sucrose in PBS, followed by a 45 min incubation in 20 % sucrose in PBS. The embryos were then left overnight in a mixture of 15 % sucrose and 7.5 % gelatin in PBS at 39 °C. The blocks were made in the same gelatin-sucrose solution the next day. Transverse cryosections (20 μm) were made on a Reichert-Jung Cryocut E cryostat and adhered to positive charged slides. On the next day, the gelatin was removed through a 30 min incubation at 39 °C in PBS, and three subsequent PBS washes at room temperature. After labeling, mounting was made using 70 % glycerol in 20 mM Tris buffer (pH 8.0). Observation was performed using a Zeiss LSM 880 laser confocal microscope, with a 63X 1.4 NA oil immersion objective.

### 2.7 Whole-mount fluorescent *in situ* hybridization

Phenylthiourea-treated embryos were fixed at 48 hpf in PFA-PBS as described, transferred to 100 % methanol on the next day, and kept at −20 °C until further use. We performed whole-mount fluorescent *in situ* hybridization (WM-FISH) using digoxigenin-labeled probes, following the protocol described by Koziol et al. (Koziol et al., 2014). The *slit2* probe was obtained from the pSPORT1-*slit2* plasmid, kindly provided by Kristen Kwan and originally generated by Yeo and collaborators (Yeo et al., 2001). Through PCR, we obtained an 897 bp probe complementary to the 3’ region of the mRNA. The sequences for the primers used are shown in Supplementary Table 1.

### 2.8 Lipophilic dye labeling

Phenylthiourea-treated embryos were fixed at 48 hpf or 5 dpf as described above. They were then immobilized on glass slides using agarose and injected with 1,1’-dioctadecyl-3,3,3′,3′-tetramethylindocarbocyanine perchlorate (DiI, Molecular Probes) or 3,3′-dioctadecyloxacarbocyanine perchlorate (DiO, Molecular Probes) dissolved in chloroform. For optic chiasm observation, DiI was injected into the vitreous chamber of one eye in 48 hpf embryos, whereas DiO was injected into the contralateral eye. For the observation of retinotopic axon distribution, DiI was injected either in the ventral or the temporal region of the retina of 48 hpf or 5 dpf larvae, at the level of the inner plexiform layer. DiO, on the other hand, was injected either in the dorsal or the nasal region of the retina, respectively. In all of the cases, after the injection the embryos or larvae were incubated for 48 hours at room temperature before mounting in agarose-glycerol as described above.

### 2.9 Time-lapse imaging

Embryos were selected around 30 hpf, anesthetized using 0.04 mg/mL MS222 (Sigma) and mounted in 1% low-melting point agarose (Sigma) over glass bottom dishes. After agarose gelification and during image acquisition, embryos were kept in Ringer’s solution (116 mM NaCl, 2.9 mM KCl, 1.8 mM CaCl2, 5 mM HEPES, pH 7.2) with 0.04 mg/mL MS222. Live acquisitions were made using a Zeiss LSM 880 laser confocal microscope, with a 40X, 1.2 NA silicone oil immersion objective. Stacks around 40 µm-thick were acquired in bidirectional scanning mode, at 1 µm spacing and 512×512 pixel resolution every 10 or 15 minutes, for 2.5-16.5 hours. The acquisition time per embryo was approximately 1 min, and up to 8 embryos were imaged in each experiment. The embryos were fixed in 4% PFA immediately after the end of the time-lapse, and processed for further confocal microscopy, labeling nuclei with methyl green.

### 2.10 Data analysis

Images were analyzed using Fiji (Schindelin et al., 2012). Optic nerve diameter was measured manually on maximum intensity z-projections of the confocal stacks. Plot profiles were built from a transect spanning the nerve, and the diameter was measured at 50 % of the peak intensity. For optic tract thickness and dorsal/ventral retinotopic segregation analysis, plot profiles were built from a transect drawn on maximum intensity z-projections from DiI/DiO-labeled samples. Volume measurements (whole retina and zn8-positive region) were performed and analyzed using intensity-based thresholding aided by manual selection as previously described (Lepanto et al., 2016). Tracking of axons to obtain distance and velocity measurements was performed using the Manual Tracking plugin (https://imagej.net/Manual_Tracking), using the site of pioneer axon crossing at the midline as reference point.

All statistical analysis was performed using GraphPad Prism software. As a routine, the data sets were checked for normality using Shapiro-Wilk normality test. In the case of normal data, we performed a Student’s *t*-test for mean comparison.

## 3 RESULTS

### 3.1 *slit2* mRNA is expressed near RGCs and their axons in the retina and proximal visual pathway

In order to assess the expression pattern of *slit2* in detail, we used whole-mount fluorescent *in situ* hybridization (WM-FISH). We focused on key stages in RGC differentiation in the zebrafish, namely 24, 32, 40, 48 and 72 hpf. No signal was detected in the retina at either 24 or 32 hpf, although a strong signal was observed along the floor plate and in different cell clusters, particularly at 30 hpf (Supplementary Fig. 1). The earliest signal in the retina was evident at 40 hpf, at the ciliary marginal zone (CMZ) and in a small group of cells at the innermost portion of the forming inner nuclear layer (Fig. 1A, B; Supplementary Video 1). Their position and general morphology indicated they might be a subset of amacrine cells. Signal at the CMZ disappeared by 48 hpf, while expression in these inner nuclear layer cells became progressively stronger from this stage (Fig. 1C, D; Supplementary Video 2) to 72 hpf (Fig. 1F, G; Supplementary Video 3). To determine the identity of these *slit2*-expressing cells, we depleted amacrine cells using a combination of two morpholino oligomers directed to the transcription factor Ptf1a, previously shown to be effective for this purpose (Fig. 1H; Almeida et al., 2014). In these embryos, *slit2* expression in the retina disappeared (Fig. 1I), while signal was still evident in other regions, such as the floor plate (Supplementary Fig. 2). Hence, the *slit2*-expressing cells in the retina are indeed a subset of amacrine cells located at the inner nuclear layer.

**Figure 1.**
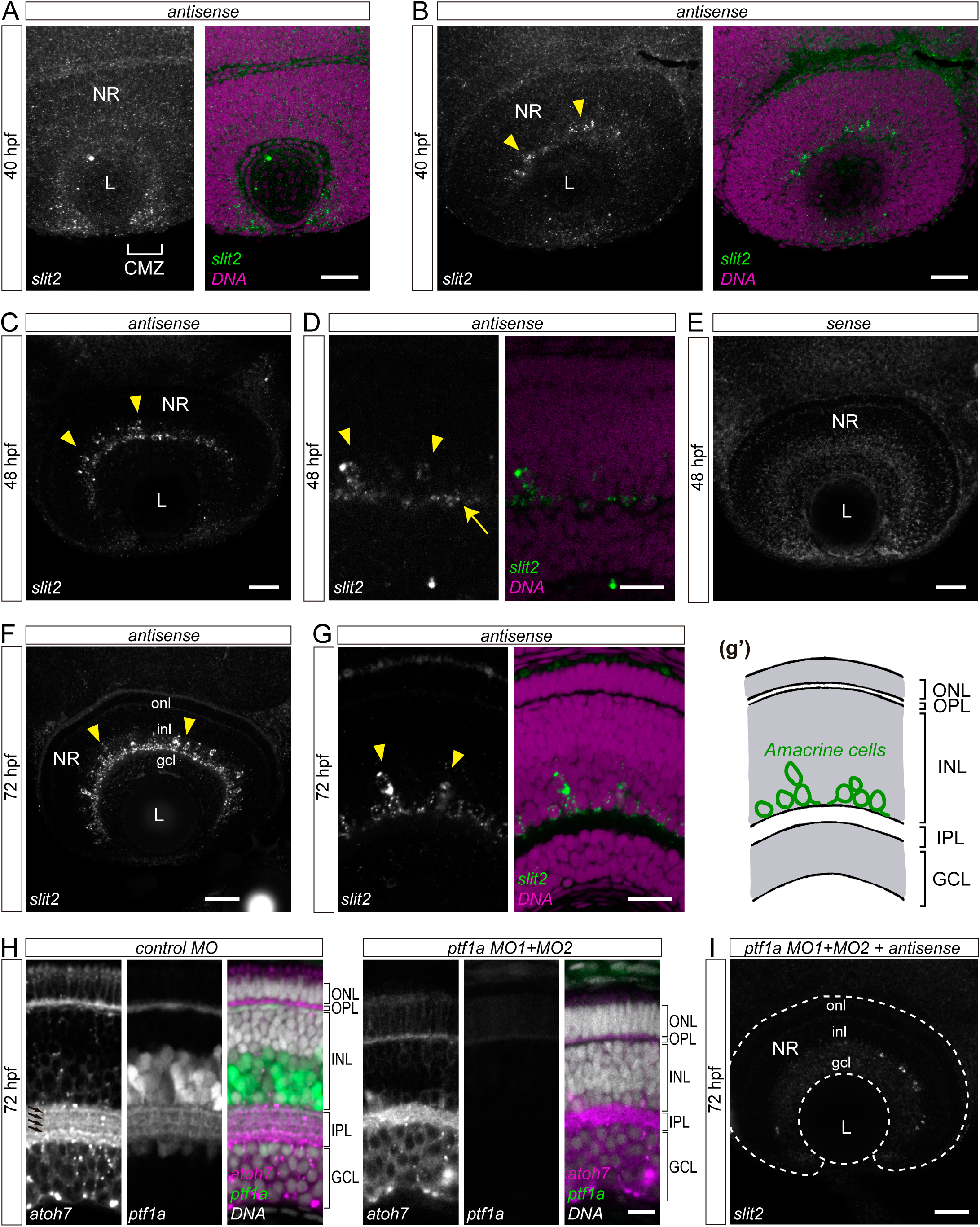
Expression of *slit2* in the developing neural retina. Whole-mount fluorescent *in situ* hybridization (WM-FISH) analysis of *slit2* in the zebrafish retina. **A, B.** Two different single confocal planes of a 40 hpf wild-type embryo. Brackets: ciliary marginal zone; arrowheads: cells located at the amacrine cell region in the central retina. See Supplementary Video 1. **C, D.** Single confocal planes of 48 hpf wild-type embryos, at different magnifications. Arrowheads: putative amacrine cell bodies; arrow: forming inner plexiform layer. See Supplementary Video 2. **E.** Control WM-FISH (sense probe) on a 48 hpf embryo. **F, G.** Single confocal planes of 72 hpf wild-type embryos. Arrowheads: putative amacrine cell bodies; g’: diagram from G highlighting these cells. See Supplementary Video 3. **H.** Single confocal plane of 72 hpf SoFa1 embryos injected with control or *ptf1a* morpholinos (MO). RGCs are labeled by *atoh7*:Gap43*-*mRFP and amacrine/horizontal cells by *ptf1a*:EGFP expression. Arrows: synaptic sublaminae. **I.** Maximum intensity z-projection of a 72 hpf wild-type embryo injected with *ptf1a* MO1+MO2, and labeled by WM-FISH to *slit2* (compare with signal in F). CMZ: ciliary marginal zone; GCL or gcl: ganglion cells layer; INL: inner nuclear layer; IPL: inner plexiform layer; L: lens; NR: neural retina; ONL or onl: outer nuclear layer; OPL or opl: outer plexiform layer. Scale bars: A-C, E, F, I, 40 μm; D, 15 μm; G,20 μm; H, 10 μm.

From 40 hpf, *slit2* expression was also evident all along the optic nerve area, inside and outside the retina (Fig. 2A; Supplementary Video 4), and by 48 hpf, expression was detected in cells surrounding the extraretinal optic nerves, as has been previously described (Fig. 2B; Supplementary Video 5; Chalasani et al., 2007), and in a few easily-individualized cells on the anterior side of the proximal optic tract (Fig. 2C; Supplementary Video 5). In addition, we observed a strong signal in two round, bilateral structures at the ventral portion of the forebrain (also described in Chalasani et al., 2007), detectable from 30 hpf, and strongly labeled by 40 hpf (Fig. 2A; Supplementary Figure 1; Supplementary Video 4). At 72 hpf, no expression was detected at the optic chiasm or the optic tectum, as previously reported (Fig. 2D, E).

**Figure 2.**
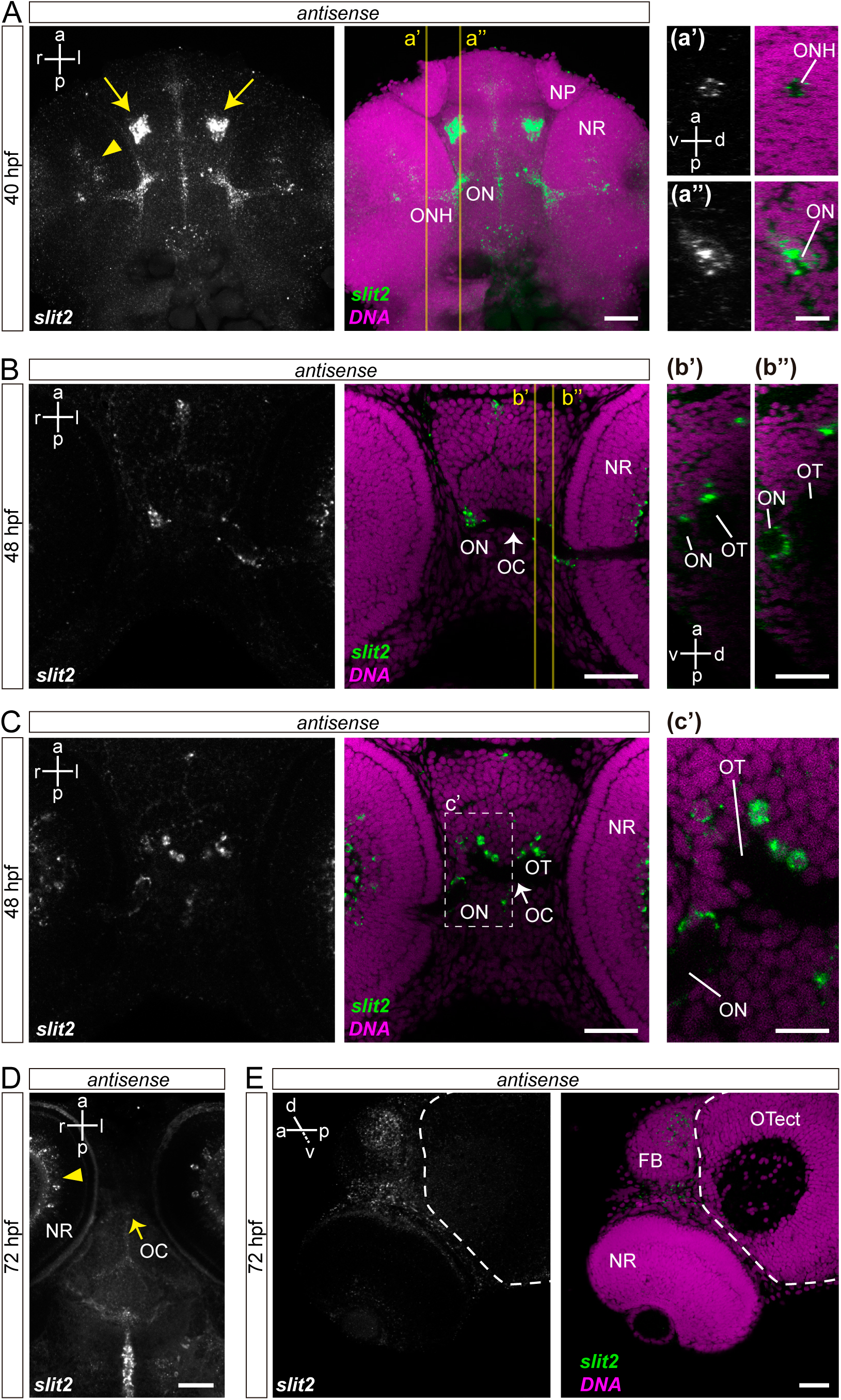
Expression of *slit2* along the pathway of RGC axons. WM-FISH analysis of *slit2* in the zebrafish brain. **A.** Maximum intensity z-projection of the cephalic region of a 40 hpf wild-type embryo (ventral view). Arrowhead: amacrine cells; arrows: cell clusters in the ventro-rostral diencephalon. a’ and a’’, magnified orthogonal sections from A. See Supplementary Video 4. **B, C.** Two different confocal planes of a 48 hpf wild-type embryo. b’ and b’’, orthogonal sections from B; c’, higher magnification from C. See Supplementary Video 5. **D.** Horizontal maximum intensity z-projection of a 72 hpf wild-type embryo. **E.** Oblique (dorso-medial to ventral-lateral) single confocal plane of a 72 hpf wild-type embryo. Dashed line: optic tectum. NP: nasal pit; NR: neural retina; OC: optic chiasm; ON: optic nerve; ONH: intra-retinal optic nerve head; OT: optic tract; OTect: optic tectum. Scale bars: A-E, b’ and b’’, 40 μm; a’, a’’, 20 μm; c’, 15 μm.

### 3.2 A zebrafish null mutant for *slit2* is viable and presents no visible defects in RGC differentiation or intraretinal axon growth and guidance

To investigate the function of Slit2 *in vivo*, we used the CRISPR/Cas9 technology to generate a zebrafish strain harboring a mutation in the *slit2* locus, as described in the Materials and Methods section. The selected mutation consists of a 10 base-pair deletion in exon 1, which causes a frameshift that leads to a protein truncation before the first LRR domain (Fig. 3A-C; Supplementary Fig. 3). As a consequence, this allele is expected to behave as a functional null. The mutation segregation, as seen from the genomic PCR analyses, is typically Mendelian (Supplementary Fig. 3). The homozygous individuals are viable and fertile, which allowed us to maintain both heterozygous and homozygous adult fish. We assessed the phenotype by incrossing either heterozygous (obtaining *slit2+/+* wild-types, *slit2+/-* heterozygous and *slit2-/-* zygotic mutant embryos) or homozygous adults (obtaining maternal-zygotic mutants, from now on denoted as *slit2-/-^mz^*). The *slit2+/-*, *slit2-/-* and *slit2-/-^mz^* embryos were externally indistinguishable from wild-type embryos, as was the case for *slit2* morpholino-treated embryos (Fig. 3D). RT-PCR analysis of early *slit2* expression revealed the presence of maternal mRNA in oocytes, with a significant reduction, albeit still detectable, by the sphere stage (4 hpf) and a very remarkable increase at 48 hpf (Fig. 3E). At this stage, maternal-zygotic *slit2-/-^mz^* mutants presented a sensibly lower amount of *slit2* mRNA (approximately 30% less than wild –type embryos), which could be caused by nonsense-mediated mRNA decay.

**Figure 3.**
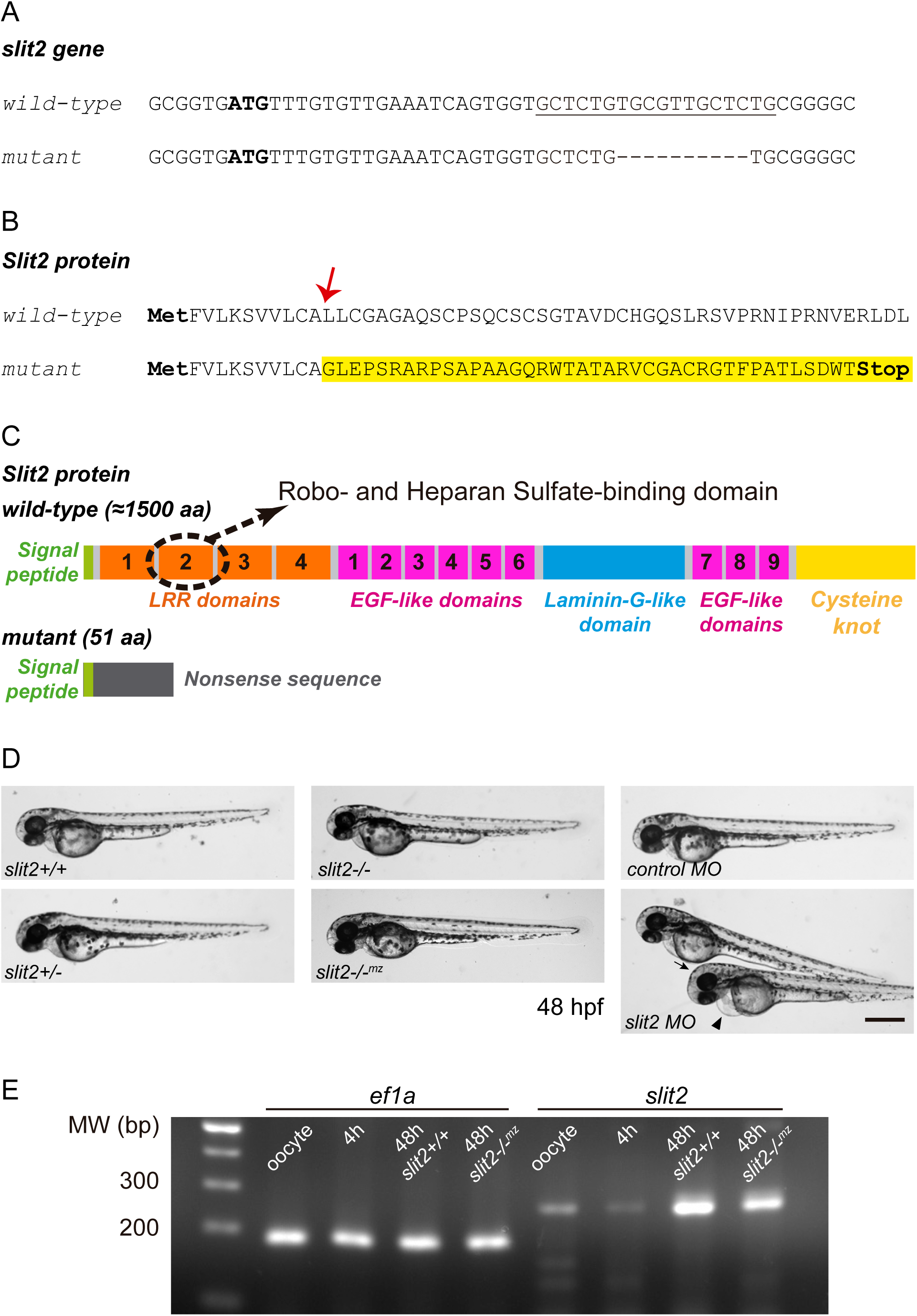
The NM_131753.1:g.30_39del mutant line presents a 10-base pair deletion in the first exon of the *slit2* gene. **A.** Initial coding sequence of the wild-type and CRISPR-Cas9-mutated *slit2* gene. The initiation ATG codon is marked in bold and the sgRNA target sequence is underlined. A 10 base-pair deletion can be observed in the mutated sequence. **B.** N-terminal sequence of the Slit2 protein. The deletion results in an mRNA coding sequence frameshift, causing a change in the protein sequence from amino acid L12 (arrow) onwards and the subsequent introduction of a premature STOP codon. **C.** Diagram representing the whole Slit2 protein (around 1500 residues long) and its main domains, compared to the resulting product upon the NM_131753.1:g.30_39del mutation. The frameshift begins at the signal peptide-encoding region and a nonsense sequence follows until the premature STOP codon, resulting in a truncated protein of 51 aa. **D.** Low magnification images of 48 hpf embryos show no evident external defects in *slit2* mutants from either heterozygous (*slit2-/-*) or homozygous (*slit2-/-^mz^*) parents. Some embryos injected with *slit2* MO exhibited mild defects such as cardiac (arrowhead) or cephalic (arrow) oedema. **E.** RT-PCR analysis of slit2 mRNA levels in wild-type oocytes, 4 hpf and 48 hpf embryos, and slit2-/-mz 48 hpf embryos. Primers to *ef1a* were used as a control. Scale bar: 500 μm.

The above-described observation that *slit2* was expressed by amacrine cells suggested it might play an intraretinal role in RGC differentiation. We set out to determine if that was the case by looking at 5 dpf larvae, where retinal lamination and RGC differentiation is mostly complete. F-actin and nuclei labeling of mutant larvae revealed that the overall structure and organization of the retina was not affected (Supplementary Fig. 4A). Moreover, the formation of sub-laminae in the inner plexiform layer did not seem to be impaired. In no case did RGC immunostaining using the anti-neurolin antibody zn8 reveal detectable defects on RGC morphology or ectopic axon growth inside the retina, as evidenced at 48 hpf (Supplementary Fig. 4B, C). There were no significant differences in the measured volume of the ganglion cell layer between wild-type and maternal-zygotic mutants (Supplementary Fig. 4D). To better assess intraretinal axon guidance and fasciculation, we digitally processed whole-eye confocal stacks of zn8-labeled 48 hpf embryos, in order to obtain planar reconstructions of the optic fiber layer as observed in Supplementary Figure 4E. We did not observe evident differences in axon bundling or directionality between control and *slit2* mutant or morpholino-injected embryos.

### 3.3 *slit2* is important for RGC axon bundling in the optic nerve

In addition to forming an apparently normal optic fiber layer, 48 hpf RGC axons neatly converged to form an optic nerve head and an optic nerve that exited the eye to meet the contralateral nerve at the optic chiasm at the embryo midline (Fig. 4A, B; Supplementary Fig. 4C). A closer look at zn8-labeled *slit2-/-^mz^* embryos revealed, however, a very conspicuous defect: their optic nerves appeared thickened (Fig. 4B).

**Figure 4.**
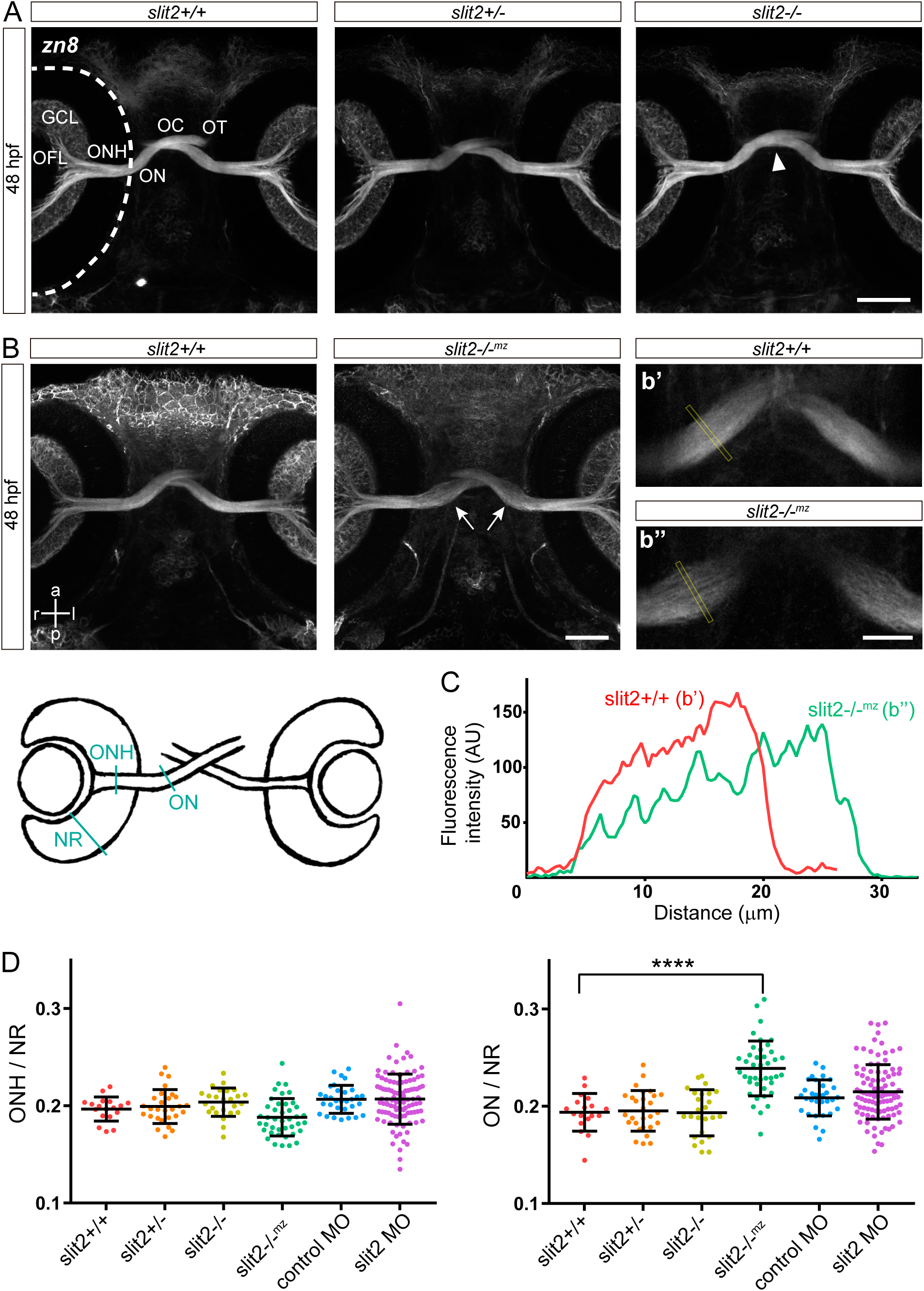
Effects of *slit2* mutation on optic nerve morphology. **A, B.** Maximum intensity z-projections of 48 hpf embryos immunostained to label RGCs (zn8 antibody). b’ and b’’ are magnified images from B, which better show the thickening of the optic nerve in *slit2-/-^mz^* embryos. Dashed line: outer retinal border. Arrowhead in A: optic chiasm defect. See Supplementary Video 6. **C**. Fluorescence intensity profile of zn8 signal in the optic nerves, measured along the transects in b’ and b’’, showing a wider and apparently more defasciculated nerve in b’’. **D.** Measurement of the optic nerve diameter, in both its intraretinal (ONH, left) and extraretinal (ON, right) portions, compared to the retina thickness. n nerves/embryos (n experiments) = 18 (3) *slit2+/+*, 28 (3) *slit2+/-*, 25 (3) *slit2-*/-, 41 (3) *slit2-/-^mz^*, 31 (3) control MO, 101 (9) *slit2* MO; mean ± SD; Student’s *t* test. The diagram shows the measured structures displayed in the graphs in D. GCL: ganglion cell layer; NR: neural retina; OC: optic chiasm; OFL: optic fiber layer; OT: optic tract. Scale bars: A, B, 50 μm; b’, b’’, 25 μm.

This phenotype was not observed in *slit2+/-* or *slit2-/-* embryos (Fig. 4A). The increase in nerve thickness was accompanied by a lower average intensity in zn8 labeling, and, at higher magnifications, it was possible to observe that axons or thin axon bundles appeared relatively separated (enlarged image b’’ in Fig. 4). These observations were further confirmed by measuring the fluorescence intensity profile across wild-type and *slit2-/-^mz^* nerves (Fig. 4C). We also quantitatively analyzed the difference in thickness, between all the conditions, for the optic nerve inside and outside the retina. No significant difference was found for intraretinal portions of the optic nerve head between mutant or morphant and wild-type embryos (Fig. 4D). However, a significantly thicker extra-retinal optic nerve was evident in *slit2-/-^mz^* embryos. We also assayed the effect of the translation-blocking *slit2* morpholino, which showed a wider data dispersion in optic nerve thickness, albeit with no significant difference to control (Fig. 4D).

Similar bundling defects were evident in a different subset of commissural axons, namely those that form the anterior commissure (AC) in the telencephalon (Supplementary Fig. 5). Axon labeling with anti-acetylated tubulin antibody revealed a thicker tract of the AC in *slit2-/-^mz^* embryos, as well as some misguided axons. This phenotype was evident from 30 hpf and became more prominent by 40 hpf. In contrast, no defects were observed in the post-optic commissure (POC) tract.

### 3.4 *slit2* is important for RGC axon organization at the optic chiasm

In several cases, zn8 labeling revealed a second defect in mutant embryos, at the level of the optic chiasm (arrowhead in Fig. 4A; Supplementary Video 6). To better characterize this observation, we differentially labeled the optic nerves using the lipophilic dyes DiI and DiO (Fig. 5). In *slit2+/+* embryos, all the axons belonging to one of the nerves crossed anteriorly (and slightly ventral) to the axons from the contralateral nerve (Fig. 5A; Supplementary Video 7), as has been previously described (Macdonald et al., 1997). In *slit2-/-* embryos, as well as in *slit2* morphants, on the other hand, one optic nerve frequently split into two groups of axons that surrounded the contralateral nerve (Fig. 5A, B; Supplementary Video 7). Surprisingly, this phenotype was observed with a higher frequency in *slit2-/-* mutant embryos than in *slit2-/-^mz^* mutants and *slit2* morphants, as assessed by zn8 labeling (Fig. 5C). Moreover, *slit2+/-* embryos also presented crossing defects, albeit with lower frequency (Fig. 5C). When considering which eye the nerve that split came from, we found that the proportion between left and right eye came close to 1:1 for all cases (Fig. 5D). This is comparable to wild-type embryos, where we found the same proportion when assessing which nerve crossed anteriorly/ventrally with respect to the other (Fig. 5D). In no case did we observe ipsilateral axonal growth. Midline crossing defects were not found in axons from another example of commissural neurons, Mauthner cells (Supplementary Fig. 6).

**Figure 5.**
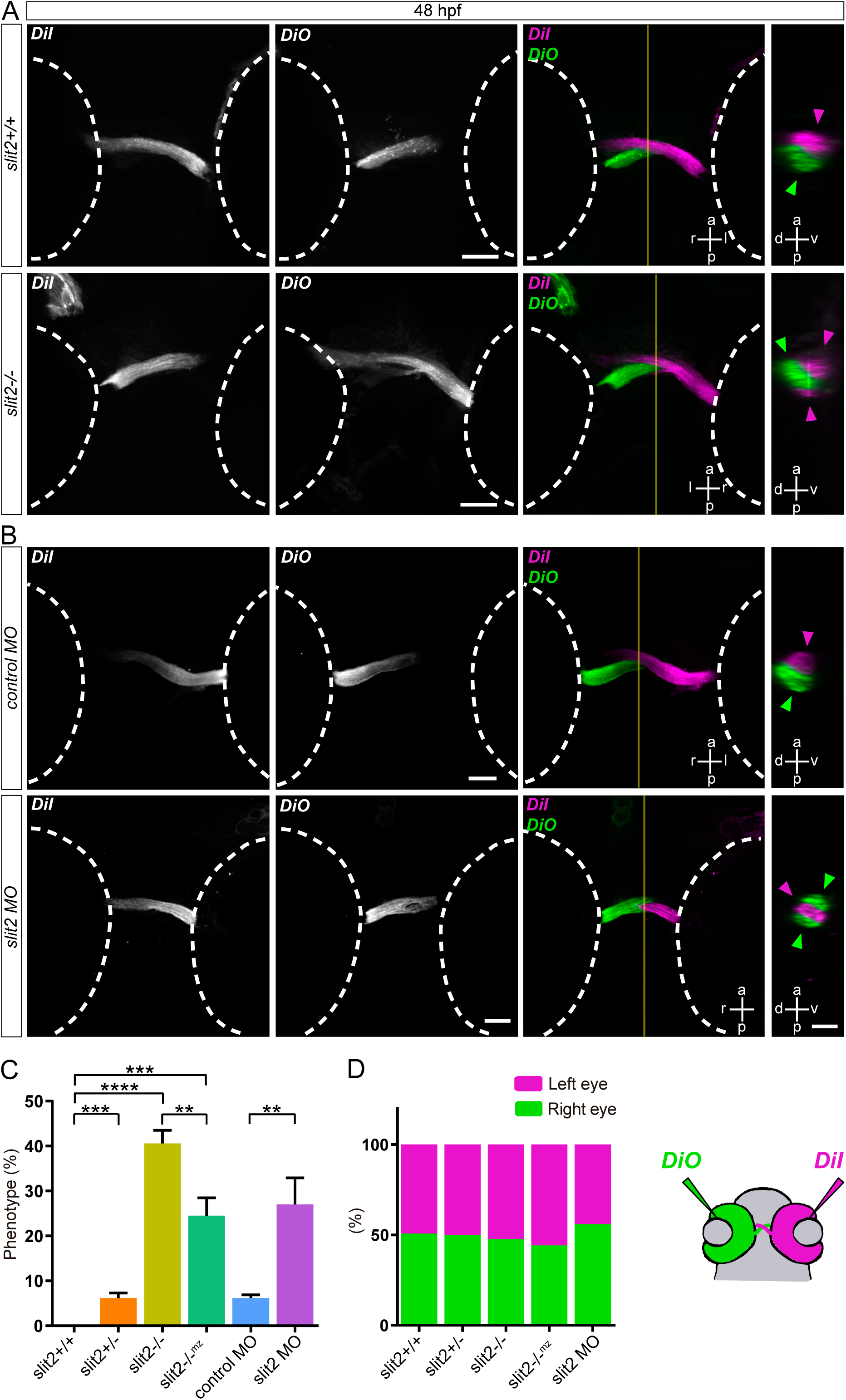
Effects of *slit2* mutation on optic chiasm formation. RGC axons from both eyes were anterogradely labeled by DiI or DiO injection into the retina. **A, B.** Horizontal maximum intensity z-projections of 48 hpf embryos, at the level of the optic chiasm, with magnified orthogonal sections at the sagittal plane level (vertical line). The maximum intensity projections showing the DiI/DiO separated channels include the proximal portion of the optic tracts, while in the merged images, the stacks are comprised only of the optic nerve and chiasm. Arrowheads: axons from one nerve cross caudal and slightly dorsal to those from the contralateral nerve in wild-type embryos; in *slit2* mutant (A) and morphant (B) embryos, one of the nerves is frequently split into two groups of axons. Dashed line: outer retinal border. See Supplementary Video 7. **C.** Quantification of phenotype penetrance in mutant and morphant embryos. n embryos (n independent experiments) = 33 (3) *slit2+/+*, 102 (3) *slit2+/-*, 50 (3) *slit2-*/-, 119 (3) *slit2-/-^mz^*, 112 (3) control MO, 140 (3) *slit2* MO; mean ± SD; Student’s *t* test. **D.** Relative frequency of left vs right optic nerve splitting in mutant and morphant embryos, compared to the frequency of left vs right nerve crossing ventrally in wild-type embryos (same embryos as in C). The diagram on the bottom right shows the strategy used for anterograde axon labeling. Scale bars: A, B, 40 μm; orthogonal sections, 20 μm.

In a further experiment, we anterogradely labeled the nasal and temporal RGCs using DiO and DiI, respectively. In control embryos, a clear segregation of nasal and temporal axons was observed along the visual pathway, where axons coming from the nasal retina always localized anteriorly to those from the temporal retina (Fig. 6). We found that this retinotopic axon sorting was partially disrupted in *slit2* MO-injected embryos at the distal optic nerve and chiasm, with axons appearing more disperse in general. When an optic nerve “splitting” phenotype was present, some axons, either from the temporal or the nasal region, surrounded the contralateral nerve at the chiasm, after separating from their own axon bundle, to then re-connect with the optic tract after crossing (Fig. 6).

**Figure 6.**
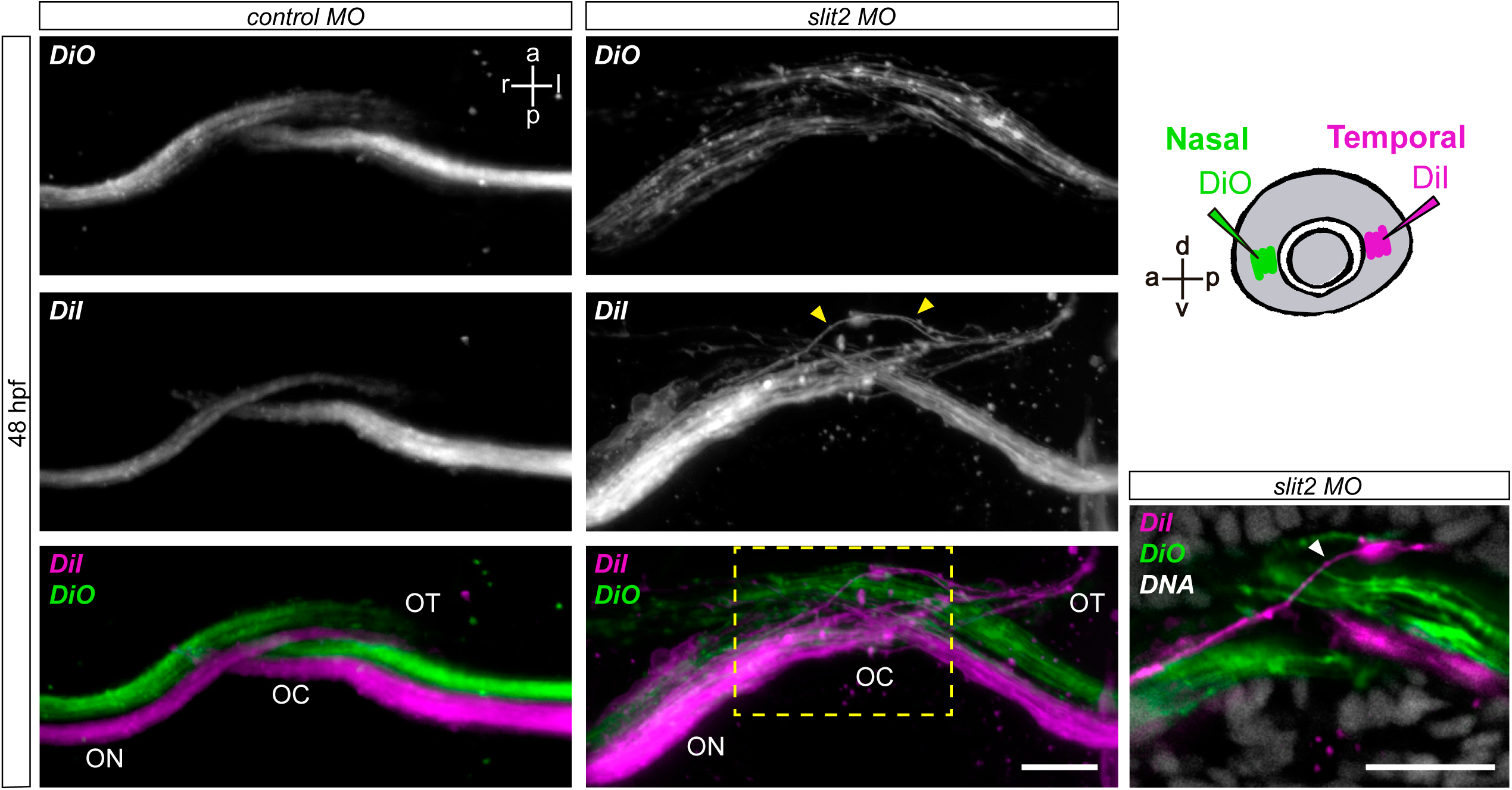
*slit2* knockdown causes axon disorganization at the optic chiasm. RGC axons from both eyes were anterogradely labeled by DiI or DiO injection into the temporal or nasal region of the retina, respectively (method illustrated in the diagram), in control or *slit2* MO-injected embryos. Maximum intensity z-projections of confocal stacks acquired from ventral are shown. Arrowheads: a small group of axons (in this case from temporal RGCs from the right eye) separate from their bundle and surround the contralateral optic nerve at the chiasm. This can be better visualized in the magnified single confocal section shown on the bottom right. Scale bars: 20 μm.

With the aim of better understanding the possible origin of this remarkable defect in optic chiasm morphogenesis, we decided to follow the growth of pioneering axons across the optic chiasm by time-lapse microscopy. We performed these experiments by injecting control or *slit2* MO in *atoh7*:Gap-EGFP transgenic embryos, in which all RGCs are evenly labeled by the expression of a membrane form of GFP (Fig. 7 and Supplementary Videos 8-10). In the control MO situation, axons usually extended quickly from the optic nerve through the chiasm, showing very little deviation from a relatively smooth curve and with small growth cones, while *slit2* morphants showed slower and more sinuous paths, with larger growth cones (Fig. 7A). These deviations resulted in defective optic chiasms, with axons crossing in a less ordered manner in all the *slit2* MO-treated embryos at the end of the time-lapse acquisition (Fig. 7B and Supplementary Videos 8-10).

**Figure 7.**
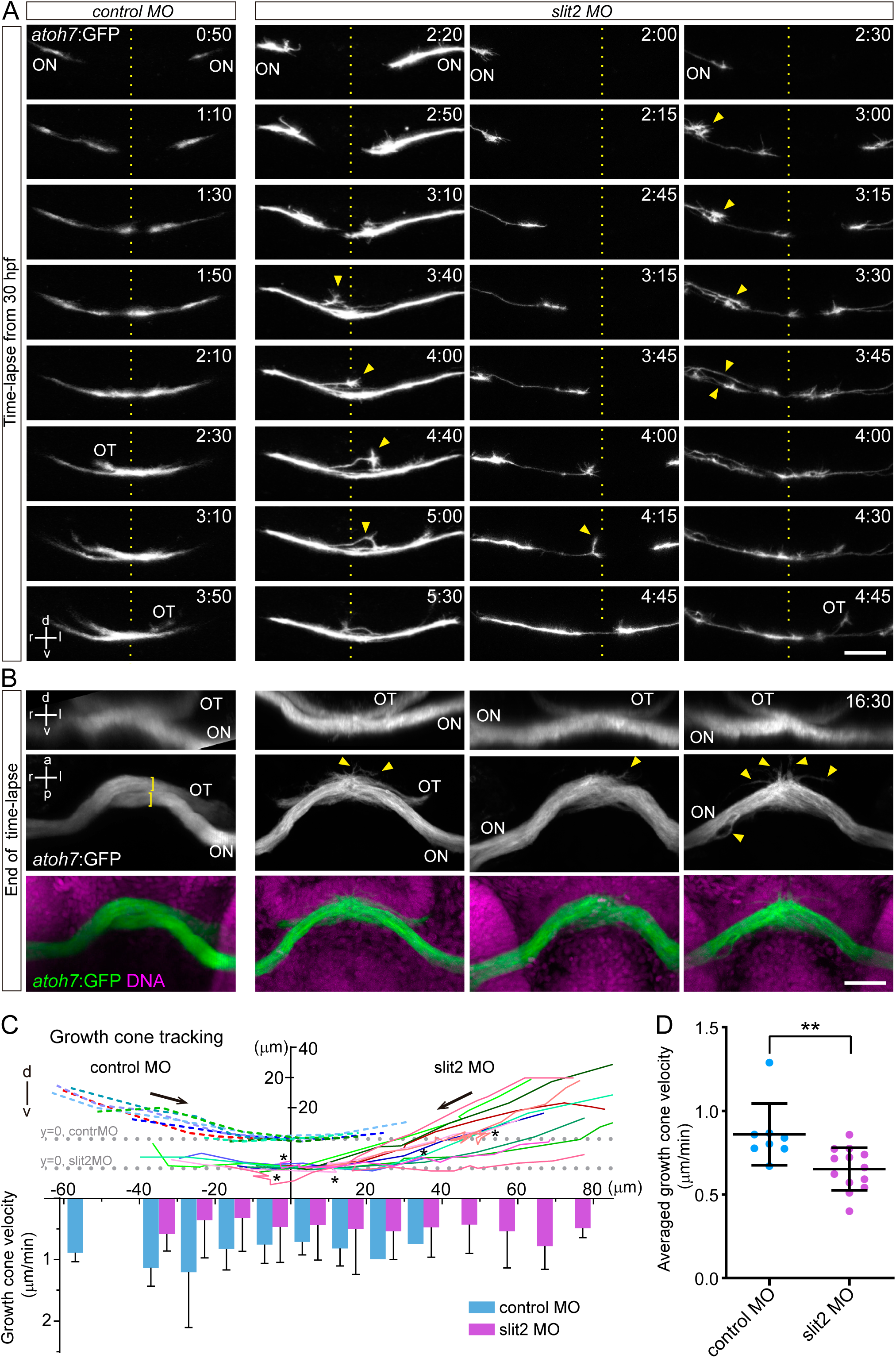
*slit2* knockdown reduces extension velocity and causes sporadic errors in axon growth around the optic chiasm. Confocal time lapse observations of pioneer optic axon growth along the area surrounding the optic chiasm, on *atoh7*: Gap43-EGFP (*atoh7*:GFP) transgenic embryos injected with control or *slit2* MO. **A.** Sequence of selected frames from time-lapse images, where the chiasm area is seen from a frontal view, and starting with the appearance of the first axon at the optic nerve. Arrowheads: axon misrouting events. See Supplementary Videos 8-10. **B.** The same embryos were fixed at the end of the time-lapse experiment (at around 46 hpf) and nuclei were labeled to evidence anatomical references. The upper row shows a frontal view and the lower ones a ventral view of the embryo’s head. Brackets: clearly segregated axon bundles from each nerve in control embryo; arrowheads: misrouted axons. See Supplementary Videos 8-10. **C.** Pioneer axon growth cones were tracked in x-y dimensions, from a frontal view. Tracks were aligned by assigning a x=0 to the brain midline and y=0 to the diencephalon ventral border. For better visualization, all control tracks are shown coming from the left side of the graph, with a 10 µm displacement in x related to the morphant ones, which come from the right side (arrows show the predominant axon growth direction in each case). The lower part of the graph shows the averaged axon growth cone velocities for 10 µm intervals along the x axis (left-right), where axon direction is the same as for the tracks; for this representation, backward growth was considered as a negative velocity value. **D.** Average of axon growth cone velocities (absolute values) all along the path observed in the time-lapse experiments, for control and *slit2* morphant embryos. Mean ± SD; Student’s *t* test. For tracking and measurements shown in C and D, 8 control MO optic nerves (from 8 different embryos) and 13 *slit2* MO optic nerves (from 11 different embryos) were analyzed in 3 independent experiments. In total, 48 (6 ± 0.9 per eye; mean ± SD) control and 181 (13.9 ± 6.7 per eye; mean ± SD) *slit2* MO time frames were considered in these analyses. Scale bars: 30 µm.

These observations were confirmed by digitally tracking the path of pioneer axons growth cones in both situations (Fig. 7C). In addition, even if not all tracked *slit2* MO axons showed sinuous paths between the optic nerve and the optic tract, their growth was significantly slower both when looking at the instantaneous velocities discriminated by their position along the path, and the averaged instantaneous velocities (Fig. 7C, D). Thus, it took axons from morphant embryos significantly longer to reach the midline (controls, 86.25 ± 45.18 min; morphants, 209.23 ± 66.98 min; mean ± SD; p<0.05 [unpaired, two-tailed Student’s *t*-test]). It is interesting to note that again, there appeared not to be a preference for axons coming either from the right or the left eye to first reach the chiasm, and neither of these aspects (laterality or crossing priority) appeared to evidently correlate with the final position of each optic nerve in controls, or bifurcation defect in *slit2* morphants.

### 3.5 *slit2* is necessary for axon sorting at the proximal optic tract

We then examined the possibility that the abnormal behavior of retinal axons when crossing the midline in *slit2* morphants resulted in alterations in axon sorting beyond the optic chiasm. For this, we again differentially labeled nasal and temporal RGC axons with the lipophilic dyes DiO and DiI, respectively, albeit in this case into a single eye, with the aim of obtaining a cleaner image of the entire visual pathway (Fig. 8 and Supplementary Videos 11 and 12). Axon segregation in control embryos was evident both at the optic nerve and the proximal optic tract, as shown by fluorescence intensity plots (Fig. 8A, C, D). This organization, however, appeared altered in *slit2* morphant embryos (Fig. 8B and Supplementary Videos 11 and 12). In this case, axon segregation at the optic nerve was only slightly affected, while the disorganization became highly conspicuous at the proximal optic tract, immediately past the optic chiasm. These defects were evidenced by the overlap of DiI and DiO curves in the fluorescence intensity plots (Fig. 8C, D). When the confocal stack was deep enough, like it was the case for the example in Figure 8, an apparent recovery of the retinotopic segregation of axons at the distal optic tract was visible (see cross section b’’’ in Fig. 8 and Supplementary Videos 11 and 12).

**Figure 8.**
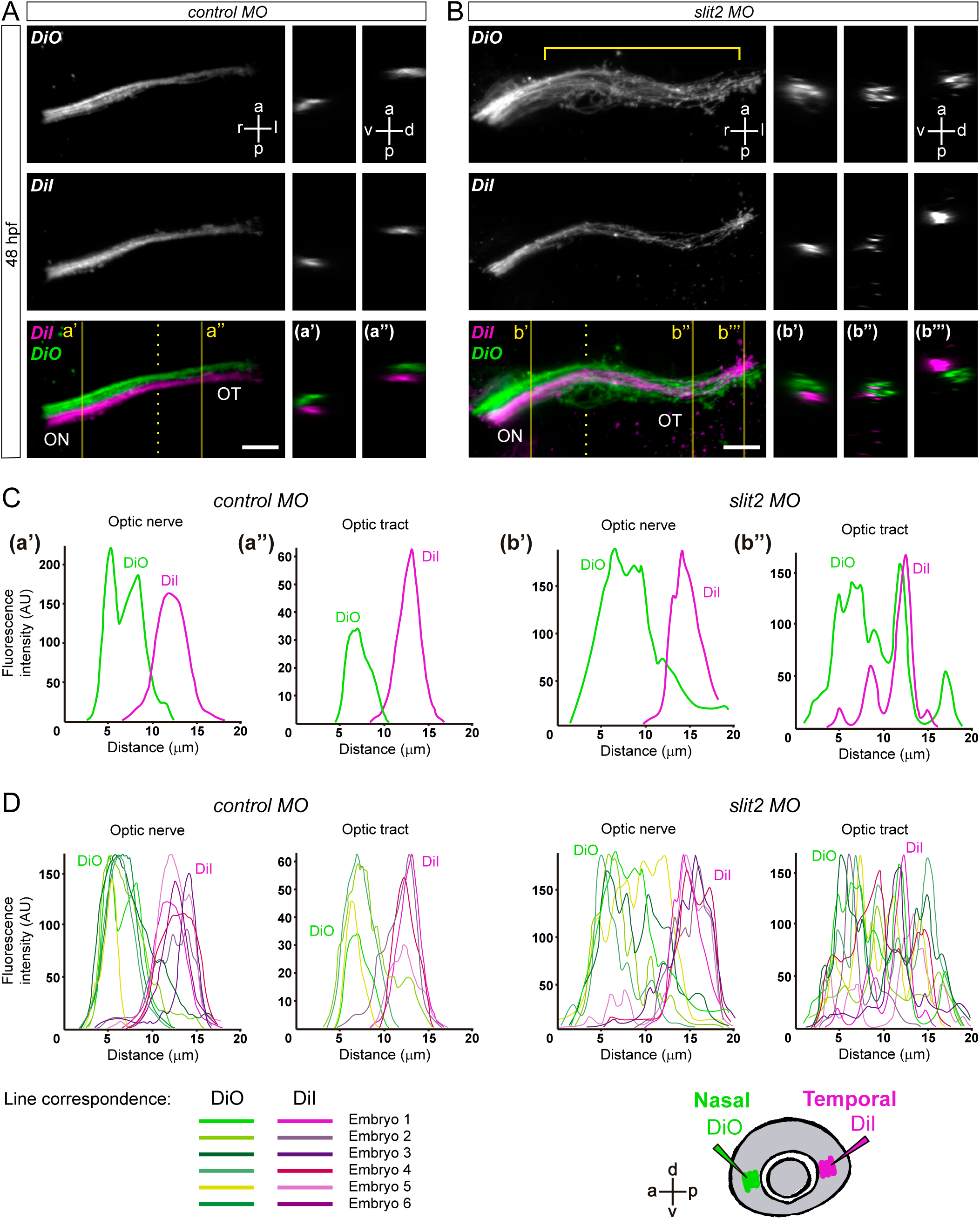
*slit2* is necessary for axon organization at the optic chiasm and proximal optic tract. Nasal and temporal RGC axons were anterogradely labeled by injection of DiO and DiI, respectively. **A, B.** Horizontal maximum intensity z-projections of 48 hpf embryos, at the level of the optic chiasm, with orthogonal sections at the sagittal plane level of the optic nerve (a’ and b’), as well as the proximal (a’’ and b’’) and distal (b’’’) optic tract. The midline is indicated by the dotted line. Bracket: extension of axon organization errors along the optic pathway of a *slit2*-MO treated embryo. See Supplementary Videos 11 and 12. **C.** Fluorescence intensity profiles through the optic nerve and optic tract of the control and morphant embryos from A and B, measured from the planes shown as orthogonal sections. **D.** Fluorescence intensity profiles through the optic nerve and optic tract of all control and morphant embryos where measurement was possible. DiI curves are shown in shades of magenta, while DiO curves are depicted in shades of green. The color code shown below indicates the correlation between the DiI and DiO curves from the same embryo. The diagram at the bottom right shows the strategy used for anterograde axon labeling. ON: optic nerve; OT: optic tract. Scale bars: 20 μm.

Given all these observations, we wondered if the lack of Slit2 would affect axon topographic sorting at the optic tectum. This was visualized in fixed whole-mount 5 dpf zebrafish larvae by injection of the lipophilic dyes DiI and DiO in the dorsal and ventral, or nasal and temporal quadrants of the retina (Fig. 9; Baier et al., 1996). After eye removal, the corresponding axonal projections from the optic tract were analyzed in lateral or dorsal view. Axon sorting at the distal end of the optic tract was not affected in *slit2-/-^mz^* larvae (Fig. 9A). Fluorescence intensity plots show that DiI- and DiO-labeled axons remained spatially separated at the distal optic tract in dorsoventral labeling (Fig. 9B). Similarly, axon sorting at the optic tectum in *slit2-/-^mz^* and *slit2* morphant embryos did not show visible defects when compared to wild-type/control embryos, in neither of the labeling strategies (Fig. 9C).

**Figure 9.**
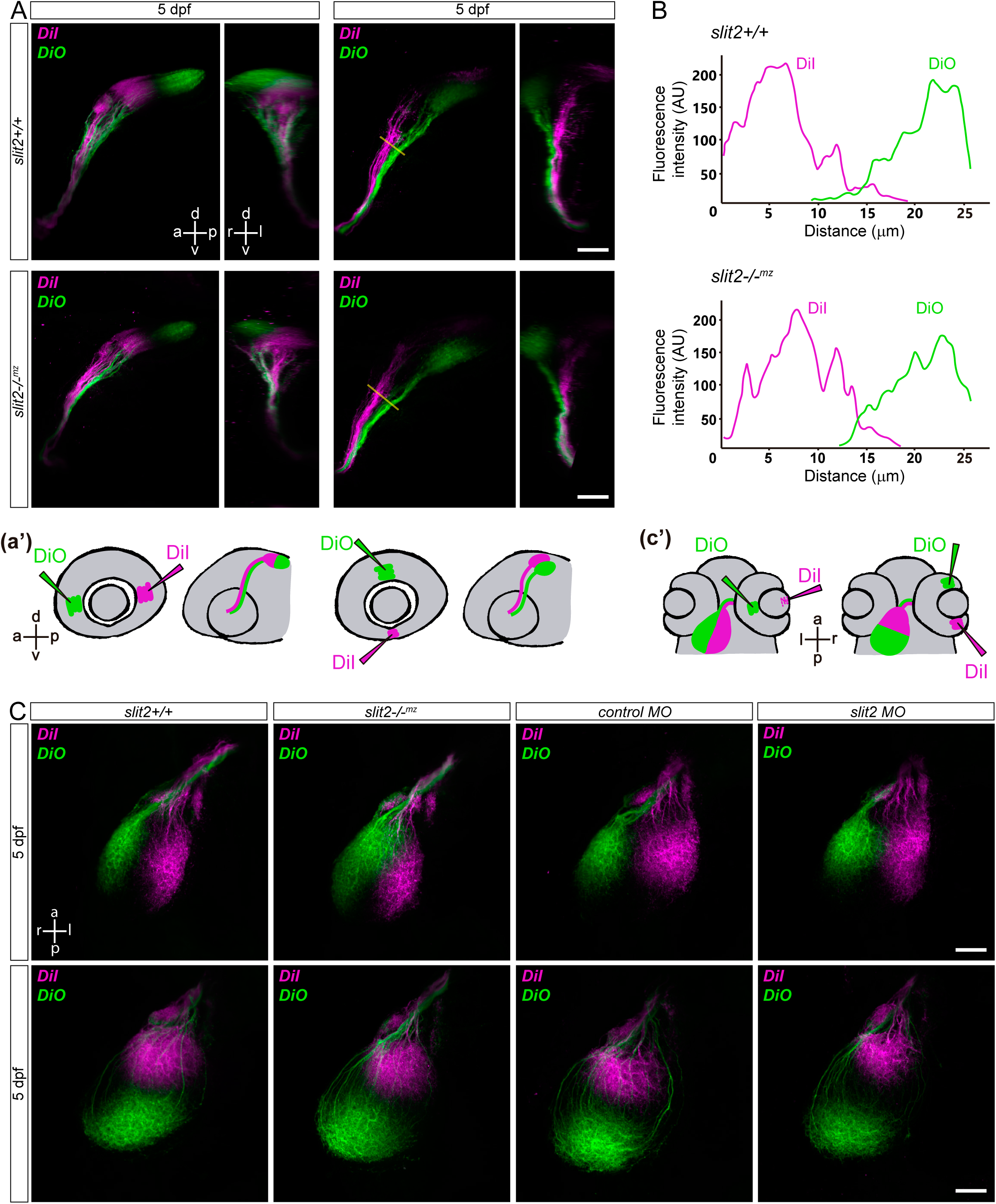
*slit2* does not play a role in axon sorting at the distal optic tract or optic tectum. Retinotopic anterograde RGC axon labeling using DiI and DiO. **A.** Parasagittal and frontal maximum intensity z-projections of the optic tract and tectal innervation of 5 dpf larvae, after either nasal/temporal or dorsal/ventral DiO and DiI injection, respectively, as depicted in the drawings (a’). **B.** Fluorescence intensity profiles through the distal optic tracts, following transects depicted as lines in A. **C.** Horizontal maximum intensity z-projections of the tectal innervation of 5 dpf larvae, in mutant and morphant situations. Injection and resulting expected labeling are depicted in c’. Scale bars: A, 40 μm; C, 30 μm.

## 4 DISCUSSION

Despite the many previous reports on the role of the Slit-Robo pathway in RGC axon growth and guidance, information on the function of Slit2 in the zebrafish RGCs has been fragmentary, partly due to the absence of a characterized mutant line. Here, we describe the generation of a null mutation in the zebrafish *slit2* locus, using CRISPR-Cas9 genome editing technology, and its primary characterization by studying the elicited phenotype on RGCs and their axons, both inside and outside the retina. A summary of the main observed effects of *slit2* mutation (and knockdown) on the visual pathway is presented in Figure 10.

**Figure 10.**
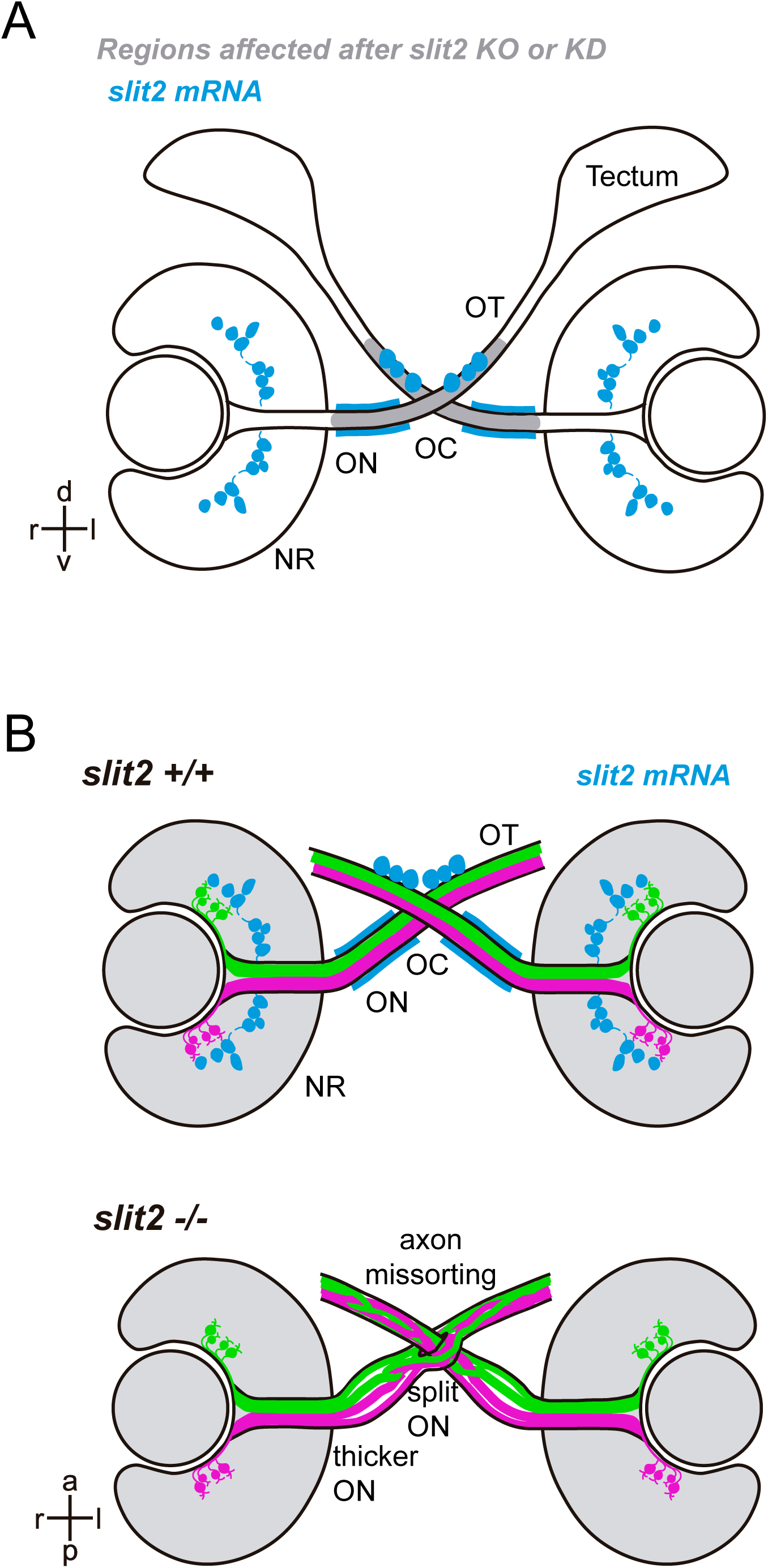
Summary of results. **A.** Frontal schematic view of a 48 hpf zebrafish embryo, illustrating the areas of *slit2* mRNA expression at that stage, and the regions exhibiting a clear phenotype upon *slit2* mutation/knockdown. **B.** Ventral schematic views of 48 hpf zebrafish embryos, highlighting the *slit2* expression areas in the wild-type, and illustrating the main phenotypic features described in the present report after *slit2* null mutation or knockdown. NR: neural retina; OC: optic chiasm; ON: optic nerve; OT: optic tract.

In the zebrafish retina, we and others (Chalasani et al., 2007) found that *slit2* was expressed by a small tier of cells at the internal part of the inner nuclear layer, and we here identified them as a subset of amacrine cells. Interestingly, a similar expression pattern was described for the mouse (Erskine et al., 2000) and *Xenopus* retina (Hocking et al., 2010; Piper et al., 2006). In all cases, displaced amacrine cells, which are localized in the ganglion cell layer, do not express detectable *slit2* mRNA levels. On one hand, this supports the idea that the polarization of the signal may be of importance and, on the other, that *slit2* expression is not determined by cell identity but by cell position. A polarized Slit2 production from the apical side of RGCs could have a function in their differentiation. Particularly, intraretinal axon guidance defects were described for mouse *slit2* mutants (Thompson et al., 2009, 2006). However, in our zebrafish mutant and morpholino analyses we did not observe obvious intraretinal defects on RGCs, neither in their morphology or number, nor in axon guidance. Furthermore, the laminar organization of the retina in these embryos appeared normal.

All the defects we observed related to RGC axons were extraretinal and localized between the optic nerve and the proximal optic tract, which very tightly correlates with *slit2* mRNA expression around the ventral midline in this region (previously also reported in the zebrafish by Chalasani et al., 2007). Our results, indicating that maternally- and zygotically-expressed Slit2 is necessary for the correct axon bundling inside the optic nerve, correlate well with previous reports that Slit2 promotes tighter fasciculation of RGC axons *in vitro* (Ringstedt et al., 2000). Interestingly, we also observed apparent bundling defects in the axon tracts that form the anterior commissure (AC). Previous studies from other researchers demonstrated *slit2* expression in the region adjacent to the AC, as well as defects in axon guidance/organization of this commissure upon *slit2* knock-down using morpholinos (Barresi et al., 2005; Hofmeister et al., 2012). On the other hand, we observed no defects at the post-optic commissure (POC), indicating some specificity on the function of Slit2 in differentially modulating axon bundling.

There are two main mechanisms that might promote axon fasciculation at the optic nerve. One of them is cell adhesion, either between axons or with surrounding glial cells (see for example Bruce et al., 2017; Masai et al., 2003). The other one is given by repulsive interactions between axons and surrounding tissues (surround repulsion), which can also favor fasciculation by channeling axons into a common path, a mechanism that has been suggested for Slits (Rasband et al., 2003). Functions in fasciculation have previously been attributed to the Slit-Robo signaling pathway in *Drosophila* (Bhat, 2017) and to Slit2 acting through Robo1 and Robo2 in mice (Jaworski and Tessier-Lavigne, 2012). The observed expression of *slit2* in cells closely surrounding the optic nerves from early stages strongly supports the idea that a channeling mechanism could be in place, delineating the path for RGC axons at the zebrafish optic nerve. Nevertheless, it should be considered that Slit-Robo signaling can also regulate cadherin-mediated cell adhesion in several systems, attenuating in some (Rhee et al., 2002; Santiago-Martínez et al., 2008) and potentiating in others (Shiau and Bronner-Fraser, 2009). As an example, Slit1b acting on nascent RGCs has a role in apical detachment through the regulation of N-cadherin in zebrafish embryos (Wong et al., 2012). Hence, an indirect effect of Slit2, modulating adhesion molecules such as N-cadherin, cannot be ruled out at this point.

A different phenotype was evident at the optic chiasm, where in a high proportion of *slit2* mutant and morphant cases one of the optic nerves appeared split in two branches that surrounded an intact contralateral nerve, instead of crossing one in front of the other as in the wild-type situation. The absence of a preference for a nerve or the other, as well as some differences evidenced between embryos, pointed to a stochastic origin of this defect. Interestingly, this “optic nerve splitting” effect was evident even in the heterozygous mutants, indicating allelic codominance. It is somehow unexpected that the frequency of errors is lower in *slit2*-/-*^mz^* than in *slit2*-/- embryos. In principle, this could be due to the fact that the maternal zygotic mutants were obtained after more crossings than the zygotic mutants, hence favoring the accumulation of gene compensation effects (El-Brolosy and Stainier, 2017).

As it has been indicated by previous research, the decision to cross the midline is probably one of the most delicately regulated steps in RGC axon growth to their targets (Herrera et al., 2019; Rasband et al., 2003). Our time-lapse imaging analyses indicate that, in the absence of Slit2, retinal axons presented larger and more complex growth cones, and frequently made mistakes, especially when approaching the midline, similar to what was reported for the *astray*/*robo2* mutant (Hutson and Chien, 2002). However, and contrary to what was observed in these mutants, the majority of the errors in the *slit2* morphants were eventually corrected, resulting in a relatively “mild” phenotype when compared to *astray* embryos (Fricke et al., 2001; Hutson and Chien, 2002). Moreover, we also found that these axons had a lower extension velocity when compared to controls, and that this decrease was maintained throughout the pathway between the optic nerve and the optic tract. This differs from what has been reported for other axon guidance cues. In Semaphorin 3d overexpressing-embryos, for example, retinal axons showed an altered morphology (larger growth cones and more filopodia) and a lower extension rate but only in the midline region (Sakai and Halloran, 2006). This suggests that these axons only become responsive to Semaphorin 3d once they approach the midline. Slit2 responsiveness, in contrast, seems to be present before, during and after midline crossing, which is consistent with its expression pattern. Surprisingly, no decrease in growth rate was found for *astray* mutants (Hutson and Chien, 2002).

We and others (Chalasani et al., 2007; Hutson and Chien, 2002) found expression of *slit2* mRNA in cells located just anterior to the optic chiasm, which could be responsible for generating a gradient channeling the axons in this area. A role for Slit2 in optic chiasm formation was also shown in mice. Plump et al. found defects in *slit1-/-;slit2-/-* double mutants (Plump et al., 2002), but no detectable phenotype in *slit2-/-* mice. However, using a different quantification approach, Down et al. reported a relatively subtle defect of the *slit2-/-* chiasm, where many axons mislocated abnormally anteriorly (Down et al., 2013). It is possible that the sorting defects we observed at the zebrafish optic chiasm come as a result of axon misguidance, a supposition that is supported by our time-lapse experiments discussed above.

Despite the chiasm phenotype observed at 48 hpf being not very severe (if compared to the *astray*/*robo2* mutant, for example), the errors made during midline crossing seem to have a notorious impact in axon organization inside the proximal optic tracts, generating errors that appear nevertheless to be corrected when axons reach the distal tract. Axon sorting at the optic chiasm has been extensively studied in mice, where it is important, for example, for ipsilateral projection. Here, dystroglycan depletion resulted in chiasm axon sorting defects (Clements and Wright, 2018), while a similar observation was made for heparan sulfate proteoglycan in the zebrafish (Lee et al., 2004). Since these extracellular matrix molecules are known to bind Slits with high affinity, it has been suggested that the axon sorting alterations occur as a result of a disruption in the extracellular distribution of Slit2 (Wright et al., 2012). On the other hand, it is important to note that a parallelism between mice and zebrafish might result difficult to establish regarding axon guidance at the optic chiasm, given some essential differences: a- in mice, which possess some degree of binocular vision, most RGC axons cross contralaterally and some take an ipsilateral turn at the optic chiasm (Jeffery and Erskine, 2005), while zebrafish lack binocular vision and present only contralateral axon crossing; b- a different set of Slit proteins are co-expressed in these regions: Slit1/2 in mice and Slit2/3 in zebrafish (Rasband et al., 2003).

When reaching the tectum, RGC axons must properly sort to innervate certain areas according to a retinotopic map (Kita et al., 2015). The absence of evident defects in distal tract organization in *slit2* morphant or mutant larvae, which also showed apparently normal naso-temporal and dorso-ventral retinotectal maps, suggest that proximal and distal optic tract axon sorting are independently regulated. This is consistent with previous reports that tract retinotectal mapping is normal in mutants in which retinal axons fail to cross the midline (Karlstrom et al., 1996; Trowe et al., 1996). It is likely that the sorting errors we observed were corrected in the distal region of the optic tract by the presence of additional guidance cues, such as Ephrin A (Gosse et al., 2008).

Furthermore, because we and others (Campbell et al., 2007) found no *slit2* expression at the optic tectum, the lack of a phenotype in this area comes as less surprising.

Altogether, our results, summarized in Figure 10, show that *slit2* mutation caused a relatively “mild” phenotype when compared to that in other axon guidance molecule mutants like *astray*. Although some evidence of the possibility of a small degree of genetic compensation was found, the similarity between the mutant and morphant phenotypes indicates that this was not a major factor. Our observations reinforce the idea that there is not one, but several, signaling molecules normally acting at the same time along the different steps in RGC axon extension, in accordance with the rich signaling molecule expression landscape that has been described in the retina and optic tectum, and all along the visual pathway (Herrera et al., 2019). In the zebrafish, these include other Slit factors such as Slit1a or Slit3, which could also act on Robo2 receptor, and partly explaining the apparent discrepancy with the stronger phenotype in *astray*/Robo2 mutants at the optic chiasm, and optic tract (Fricke et al., 2001; Hutson and Chien, 2002). Nevertheless, we provide here conclusive evidence for an essential function of Slit2 in the organization of the medio-ventral portion of the visual pathway of the zebrafish, comprising the optic nerve, the optic chiasm and the proximal optic tract.

## ABBREVIATIONS

CRISPR: clustered regularly interspaced short palindromic repeats
MO: morpholino oligomer
RGC: retinal ganglion cell

## Acknowledgements

The authors thank William A. Harris, University of Cambridge, and José L. Badano, Institut Pasteur Montevideo, for continued support with lab space and materials, as well as fruitful discussion; Uriel Koziol and Matías Preza, Biología Celular, Facultad de Ciencias, UdelaR, for help with whole-mount fluorescent *in situ* hybridization; Gabriela Bedó and Ileana Sosa, Genética, Facultad de Ciencias, for help with RT-PCR; Juan Pablo Fernández, Yale University, for invaluable help and discussion on CRISPR-Cas9 technique; Kristen Kwan for providing plasmids; Casandra Carrillo and Gisell González, Zebrafish Lab, Institut Pasteur Montevideo, for fish maintenance and care; Marcela Díaz and Tabaré De Los Campos, Microscopy Unit, Institut Pasteur Montevideo, for technical assistance on microscopy.

## Supplementary material legends

**Supplementary Figure 1.**
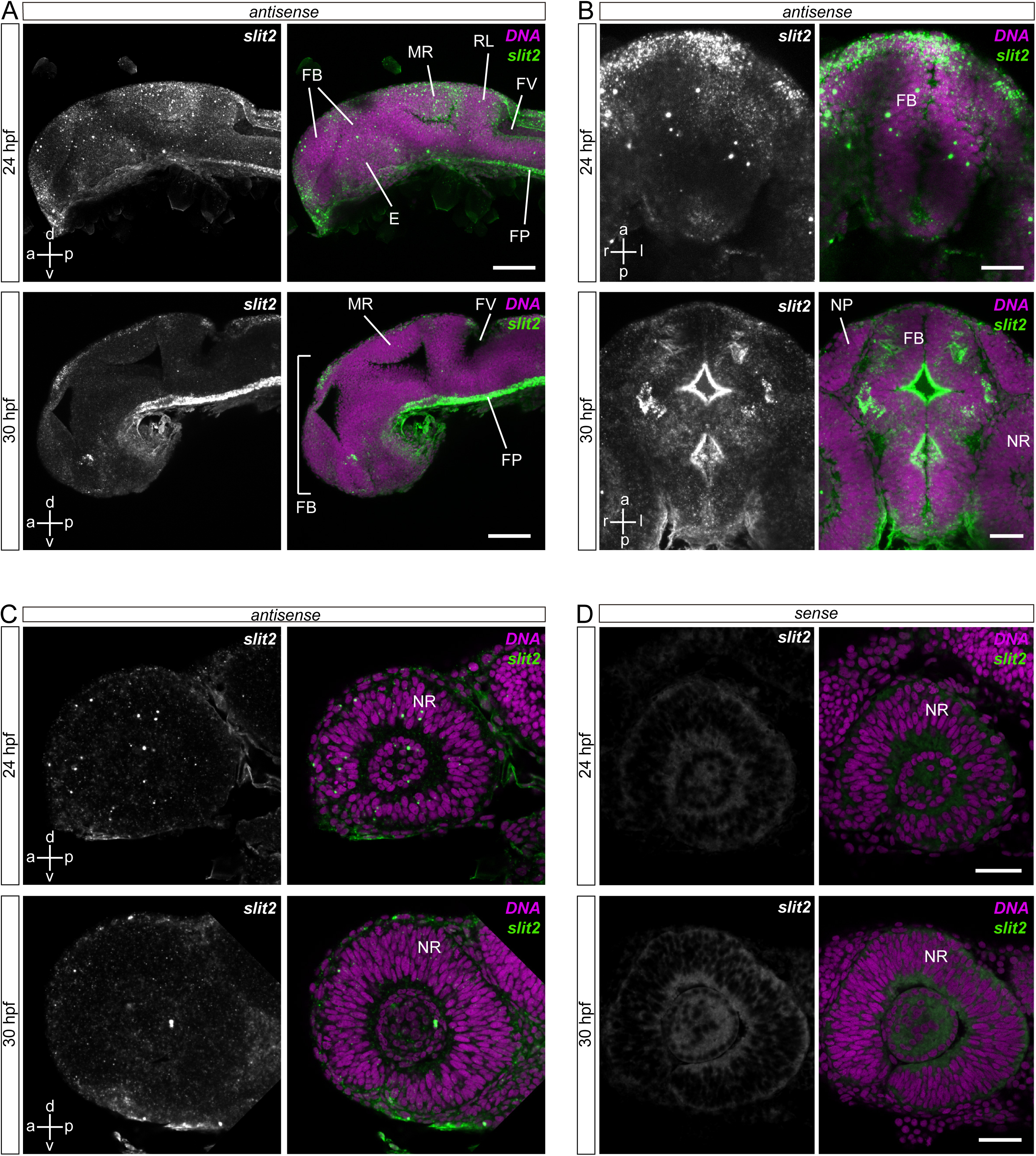
*slit2* expression at early developmental stages. Expression of *slit2* in the cephalic region of 24 and 30 hpf zebrafish embryos, evidenced through WM-FISH and confocal imaging. **A.** Parasagittal maximum intensity z-projection of the anterior region (including forebrain, midbrain and hindbrain) of 24 and 30 hpf embryos. A strong signal was observed along the floor plate (FP) and, much fainter, on the mesencephalic roof (MR), only at 24 hpf. **B.** Horizontal maximum intensity z-projection of the cephalic region (including forebrain, midbrain and eyes) of 24 and 30 hpf embryos, where *slit2* signal can be observed in different cell clusters, particularly at 30 hpf (the strong fluorescence on the ventricular surface is most probably non-specific). **C.** Parasagittal single confocal section of the eye, showing no detectable expression in the retina at 24 or 30 hpf. **D.** Confocal section from control embryos (sense probe). E: eye; FB: forebrain; FP: floor plate; FV: fourth ventricle; MR: mesencephalic roof; NP: nasal pit; NR: neural retina; RL: rhombic lip. Scale bars: A, 80 μm; B, 30 μm; C, D, 40 μm.

**Supplementary Figure 2.**
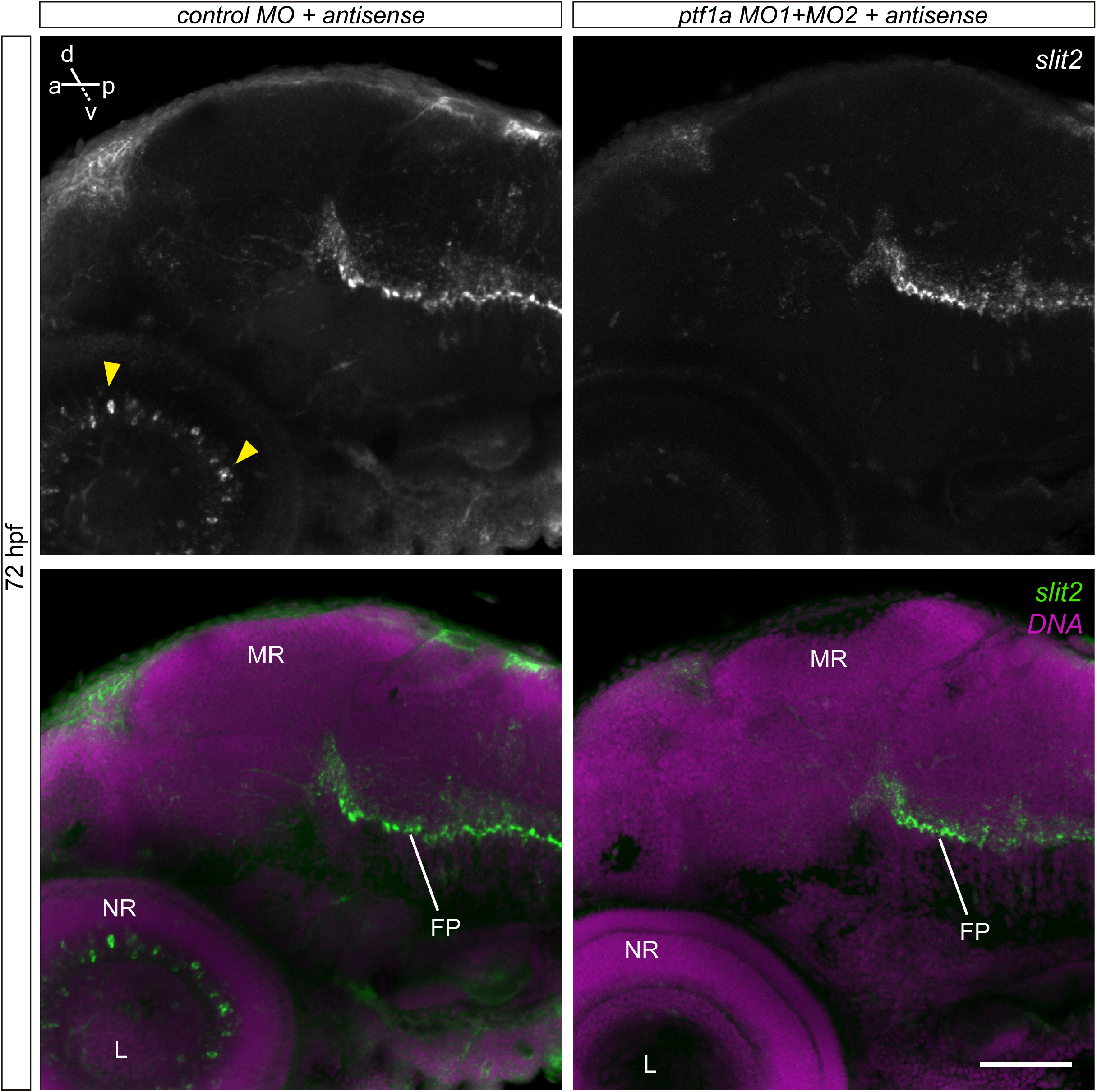
Expression of *slit2* in the retina is restricted to amacrine cells. Maximum intensity projections from 72 hpf embryos labeled by WM-FISH to *slit2*, and injected with control or *ptf1a* morpholino oligomers. In morphant embryos, *slit2* signal disappears selectively in amacrine cells of the retina (arrowheads), while it is maintained at the floorplate. FP: floor plate; L: lens; MR: mesencephalic roof; NR: neural retina. Scale bar: 60 μm.

**Supplementary Figure 3.**
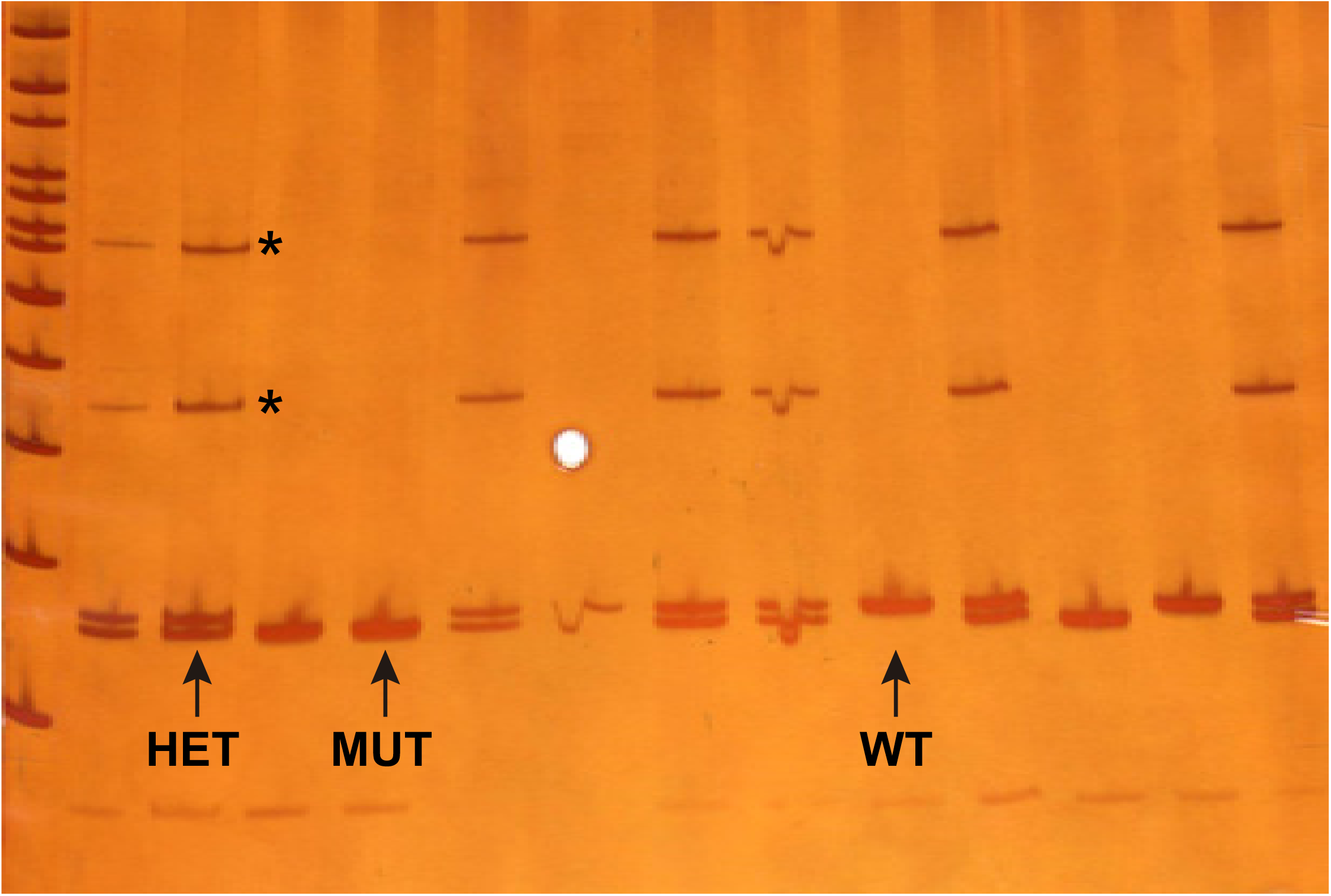
The NM_131753.1:g.30_39del mutant line can be identified by gel electrophoresis. Polyacrylamide gel electrophoresis of genomic PCR amplification products surrounding the sgRNA target sequence, from wild-type (WT), heterozygous mutant (HET) and homozygous mutant (MUT) individuals. The occurrence of the mutation can be readily identified, making it possible to differentiate the three genotypes. Heteroduplex formation in amplicons from heterozygous fish can be observed as characteristic lagging bands (asterisks).

**Supplementary Figure 4.**
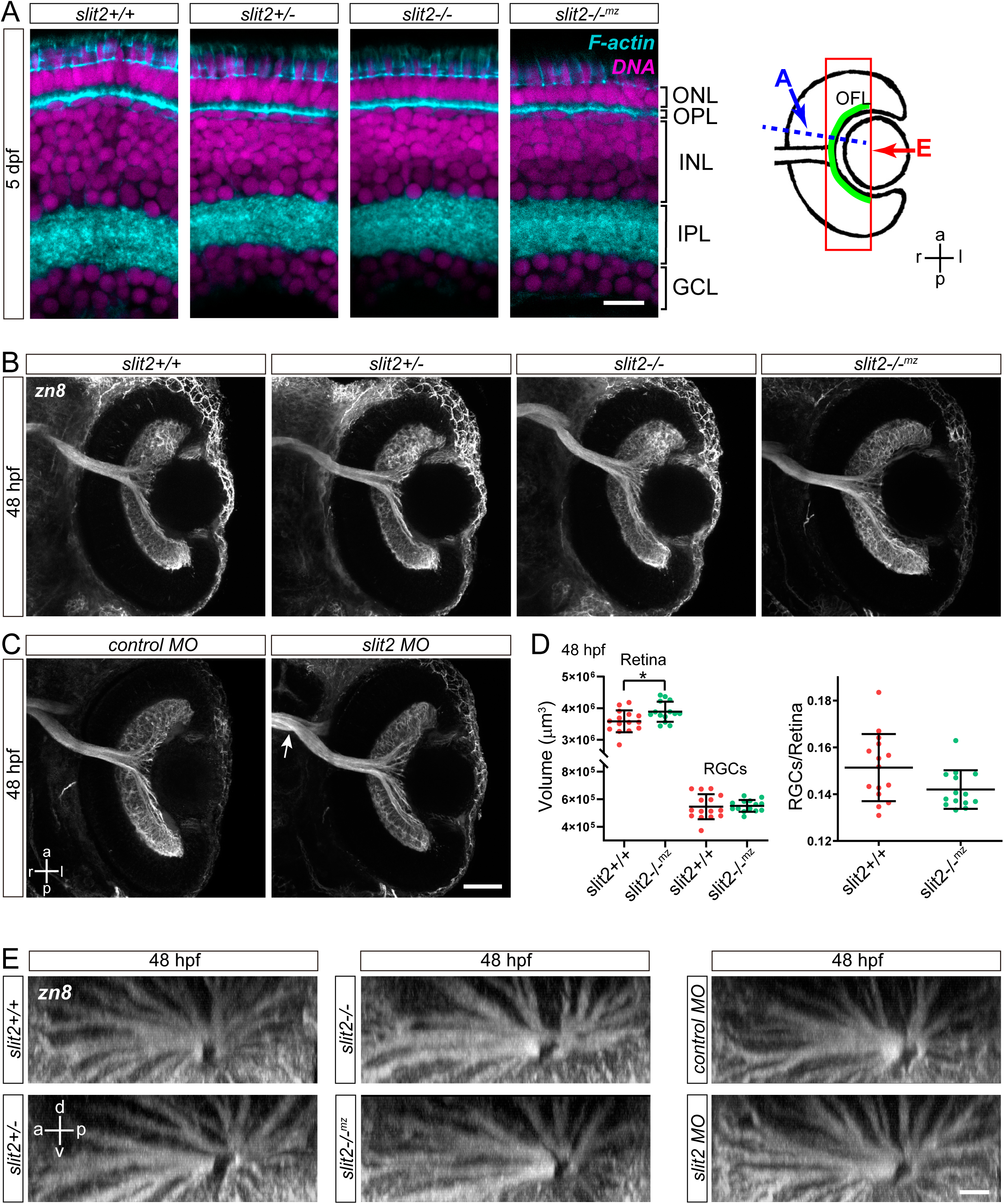
Absence of visible intraretinal axon guidance or RGC layer formation defects in *slit2* mutants or morphants. **A.** Single confocal planes from cryosections of 5 dpf larvae, labeled with phalloidin-rhodamine and methyl green. Neither the retina laminar organization nor the inner plexiform layer sub-lamination are visibly affected in mutant larvae. The diagram at the right depicts an eye section like in B-C, illustrating the regions and angles of image acquisition in A and E. **B, C.** Horizontal maximum intensity z-projections of the retina of 48 hpf embryos after immunolabeling RGCs with zn8 antibody. No RGC axons were seen growing inside the neural retina in neither morphant nor mutant embryos. The crossing defects at the optic chiasm were evident in some embryos (arrow). **D.** Quantification of the retinal and ganglion cell layer volumes in 48 hpf *slit2+/+* and *slit2-/-^mz^* embryos labeled as in A. Although there is a mild apparent increase in total retinal volume in mutants, the statistical significance is very low (p=0.02), and no significant differences were determined for the ratio RGCs/Retina. n embryos (n experiments) = 15 (2) *slit2+/+*, 14 (2) *slit2-/-^mz^*. Mean ± SD. Student’s *t* test. **E.** Lateral view from 3D-projections of 48 hpf retinas showing the organization of the optic fiber layer. No differences are observed between mutant or morphant embryos and controls. GCL: ganglion cell layer; INL: inner nuclear layer; IPL: inner plexiform layer; ONL: outer nuclear layer; OPL: outer plexiform layer. Scale bar: A, 10 μm; C and E, 40 μm.

**Supplementary Figure 5.**
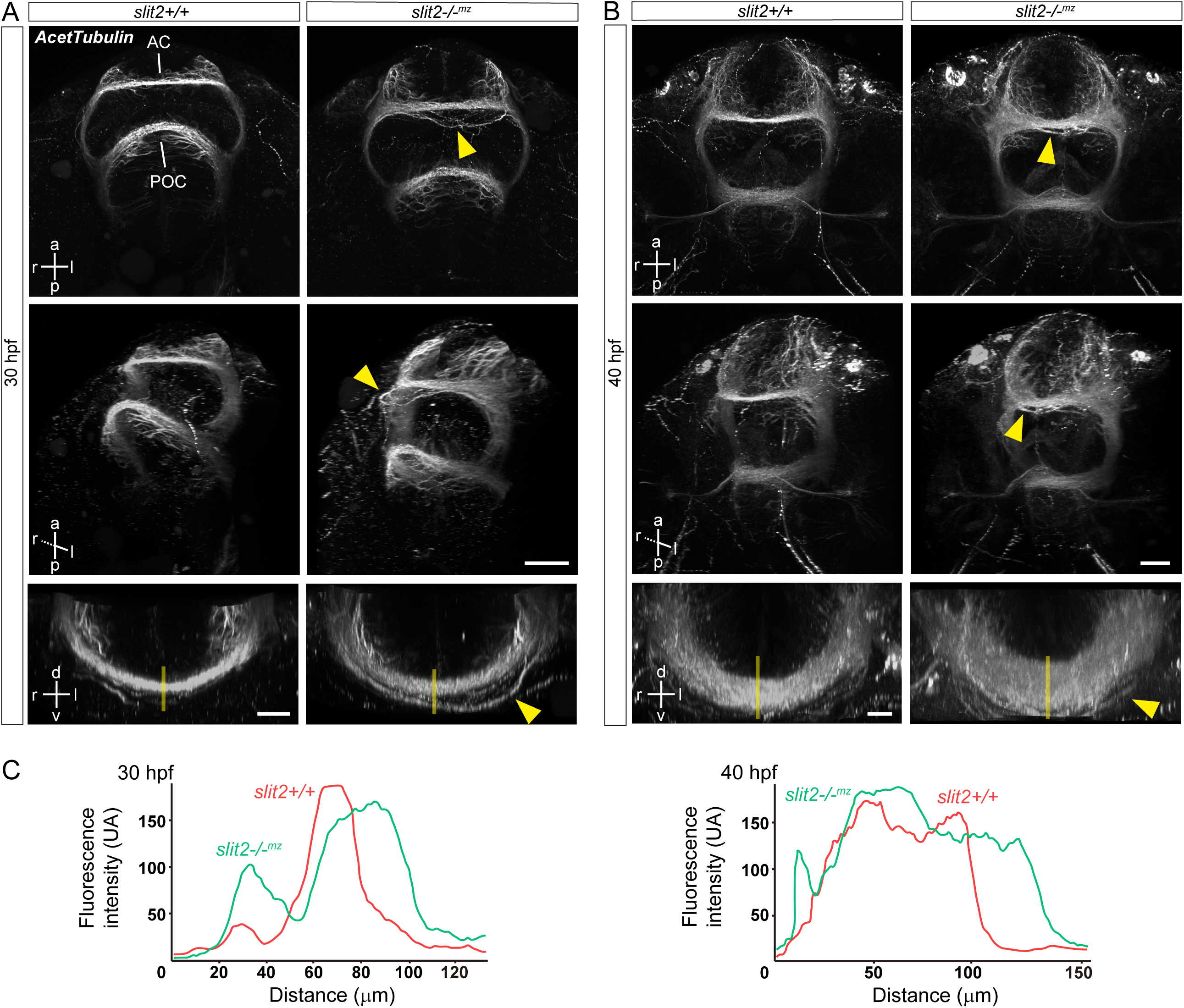
*slit2* is important for anterior commissure axon fasciculation and guidance. Immunolabeling of axons with anti-acetylated α-tubulin antibody on 30 and 40 hpf embryos. **A.** Maximum intensity z-projection of the cephalic region of 30 hpf *slit2+/+* and *slit2-/-^mz^* embryos (upper row: ventral view). Axons from the nuclei of the tract of the anterior and post-optic commissures can be seen crossing the midline, forming the anterior commissure (AC) and the post-optic commissure (POC). This can be further visualized in the 3D-projections shown in the middle row, where misguided axons can be seen along the AC in the mutant embryo (arrowheads). The axons that form the AC are tightly bundled in the wild-type and partially defasciculated in the mutant, as is visible in the magnified frontal view of the lower row. **B.** Similar images to B, from 40 hpf embryos. By this stage, more axons have incorporated into both the AC and the POC. The AC is notoriously wider in mutant than in wild-type embryos, as is clearly visible in the magnified frontal view in the lower row. **C.** Fluorescence intensity profiles through the AC, following transects depicted as lines in the AC images in the lower row in A and B. Scale bars: upper and middle row, 40 μm; lower row, 60 μm.

**Supplementary Figure 6.**
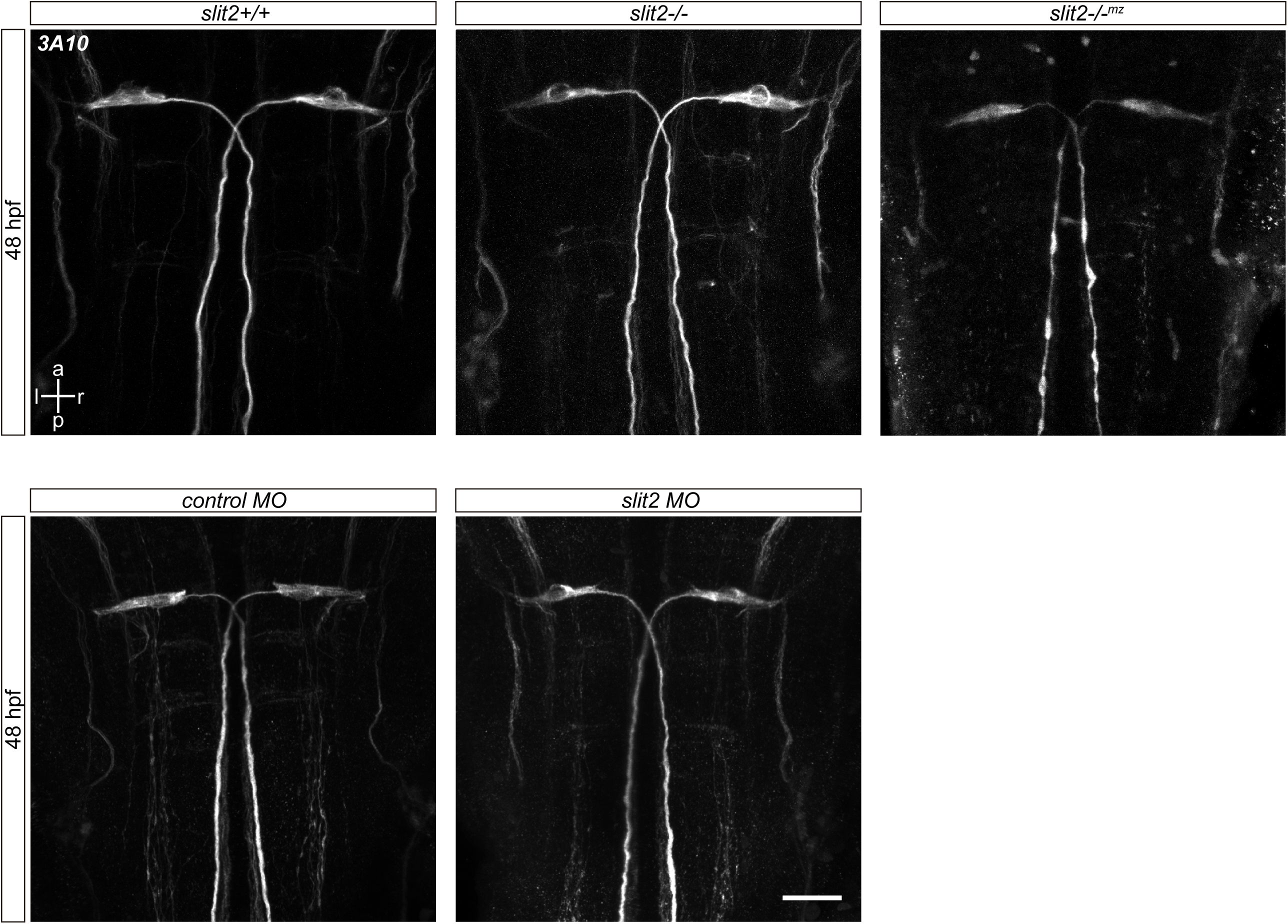
*slit2* is not essential for midline crossing of Mauthner cell axons. Immunolabeling of Mauthner cells with 3A10 antibody on 48 hpf embryos. No axon guidance defects are observed in these cells, in neither mutant nor morphant embryos. Scale bar: 30 μm.

**Supplementary Video 1. *slit2* is expressed in the ciliary margin zone and putative amacrine cells at 40 hpf.** 3D projection of a z-stack of a 40 hpf retina after WM-FISH for *slit2*. Signal can be observed at the ciliary margin zone, as well as in cells positioned in the middle region of the retina, which will probably give rise to amacrine cells. Corresponding to Fig. 1A, B.

**Supplementary Video 2. *slit2* is expressed by amacrine cells at 48 hpf.** 3D projection of a z-stack of a 48 hpf retina after WM-FISH for *slit2*. By this stage, signal is no longer detected at the CMZ, while the expression by amacrine cells becomes stronger. Corresponding to Fig. 1C.

**Supplementary Video 3. *slit2* is expressed by amacrine cells at 72 hpf.** 3D projection of z-stack of a 72 hpf retina after WM-FISH for *slit2*. By this stage, amacrine cells are the only *slit2*-expressing cells in the retina. Corresponding to Fig. 1F.

**Supplementary Video 4. *slit2* is expressed around the optic nerve area at 40 hpf.** 3D projection of a z-stack of the cephalic region of a 40 hpf embryo after fluorescent *in situ* hybridization WM-FISH for *slit2*. Signal can be detected around the optic nerve both inside and outside the retina, as well as in two bilateral structures located in the forebrain. Corresponding to Fig. 2A.

**Supplementary Video 5. *slit2* is expressed in cells surrounding the optic nerve at 48 hpf.** 3D projection a z-stack of the cephalic region of a 48 hpf embryo after WM-FISH for *slit2*. Signal can be detected in cells located around the optic nerves, as well as in a few cells just anterior to the optic chiasm and tract. Corresponding to Fig. 2B, C.

**Supplementary Video 6. *slit2* is important for optic nerve axon segregation at the optic chiasm I.** Z-stacks showing the optic chiasm of *slit2+/+* and *slit2-/-* embryos at 48 hpf, after immunostaining with zn8 antibody. In *slit2+/+* embryos, one optic nerve can be seen crossing anteriorly to the contralateral nerve, while in *slit2-/-* one of the optic nerves splits and surrounds the contralateral nerve. The stack sequence is shown from the ventral to the dorsal region of the embryo. Corresponding to Fig. 4A.

**Supplementary Video 7. *slit2* is important for optic nerve axon segregation at the optic chiasm II.** 3D projections of z-stacks of the optic chiasm of wild-type embryos injected with control or *slit2* MO and observed at 48 hpf, where RGC axons from both eyes were labeled anterogradely with either DiI or DiO. In control embryos, one of the optic nerves can be seen crossing anteriorly and slightly ventral to the contralateral nerve, while remaining physically separate. In *slit2* morphants, one of the optic nerves can be seen splitting into two groups of axons which surround the contralateral nerve. Corresponding to Fig. 5B.

**Supplementary Video 8. Axon growth cone dynamics is altered in *slit2* MO-injected embryos.** Confocal time-lapse (4D) maximum intensity projection images of the optic chiasm region from *atoh7*:Gap-GFP transgenic embryos injected with control or *slit2* MO, acquired every 10 min from 30 hpf. At the end, a full stack reconstruction of the methyl green-counterstained embryos 16.5 h after the beginning of the time-lapse experiments is shown. In the *slit2* MO-injected embryos, axons tend to exhibit larger growth cones and some of them suffer subtle deviations from their path as they approach and cross the optic chiasm area. At the end of the experiment, these optic chiasms show clear defects in the sorting of axons coming from each eye. Corresponding to Fig. 7, also see Supplementary Videos 9 and 10.

**Supplementary Videos 9 and 10. Axon growth cone dynamics is altered in *slit2* MO-injected embryos: two further examples.** Confocal time-lapse (4D) maximum intensity projection images of the optic chiasm region from *atoh7*:Gap-GFP transgenic embryos injected with *slit2* MO, acquired every 15 min from 30 hpf. At the end, a full stack reconstruction of the methyl green-counterstained embryos 16.5 h after the beginning of the time-lapse experiments is shown. Corresponding to Fig. 7, also see Supplementary Video 8.

**Supplementary Video 11. *slit2* is important for axon sorting at the optic chiasm and proximal optic tract.** 3D projection of z-stacks of the optic chiasm of embryos injected with control or *slit2* MO and observed at 48 hpf, where axons from nasal and temporal RGCs were labeled anterogradely with DiO or DiI, respectively. In control embryos, both populations of axons remain segregated at the optic nerve and proximal optic tract, with nasal axons crossing anteriorly to temporal axons. In *slit2* morphants, this organization is mostly maintained at the optic nerve, but becomes significantly altered past the optic chiasm. Corresponding to Fig. 8, also see Supplementary Video 12.

**Supplementary Video 12. *slit2* is important for axon sorting at the optic chiasm and proximal optic tract.** Animation of transverse virtual sections along the visual pathway of control and slit2 MO-injected embryos at 48 hpf, from the proximal optic nerve to the beginning of the distal optic tract. Axons from nasal and temporal RGCs were labeled anterogradely with DiO or DiI, respectively. The gradient in axons sorting defects in the slit2 morphant start to be visible at the distal portion of the optic nerve, to become progressively more severe through the optic chiasm and at the proximal optic tract. At the distal tract, the axons regain their retinotopic segregation pattern. Corresponding to Fig. 8, also see Supplementary Video 11.

## SUPPLEMENTARY TABLES

**Table 1.**
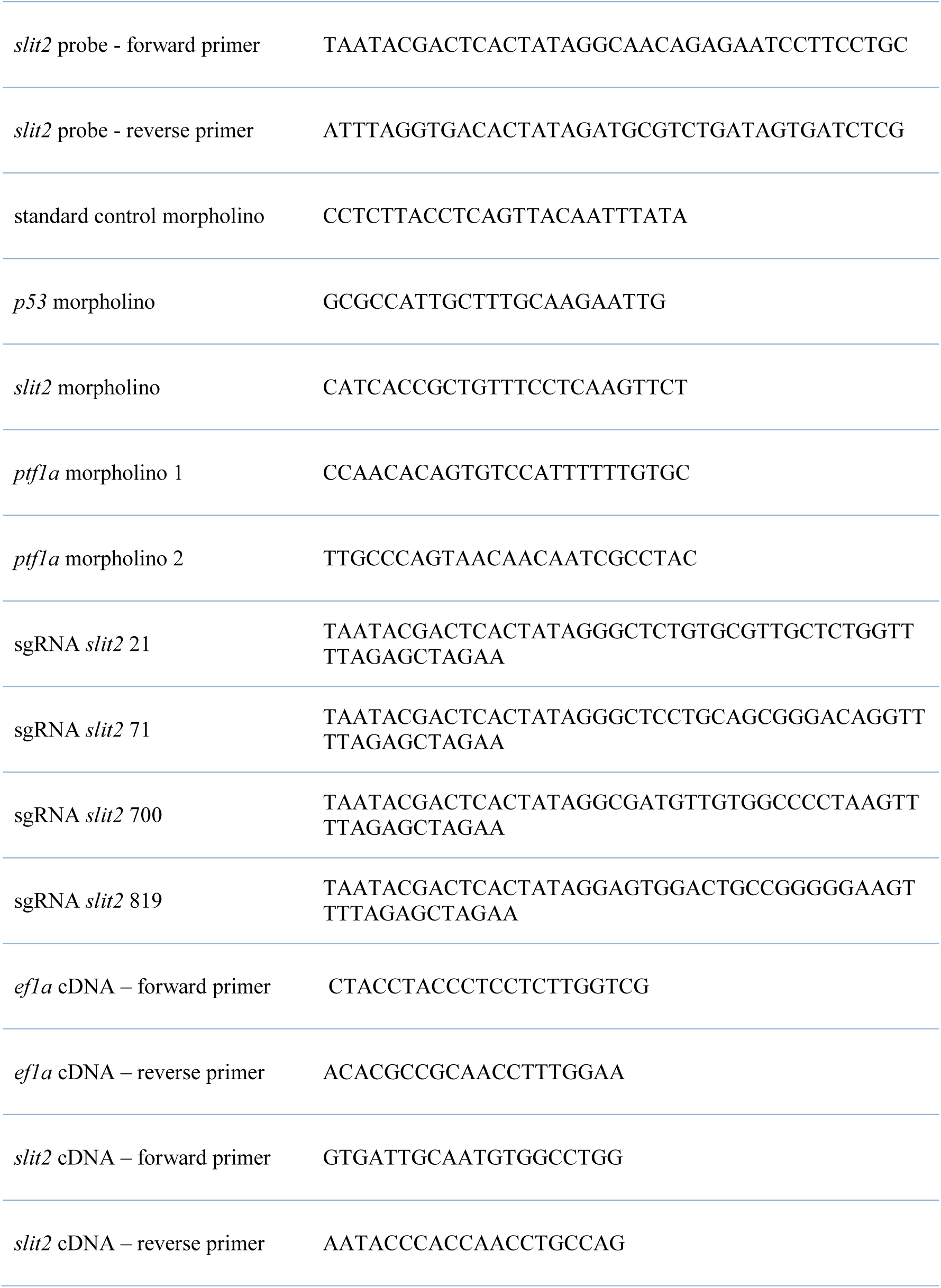

**Table 2.**
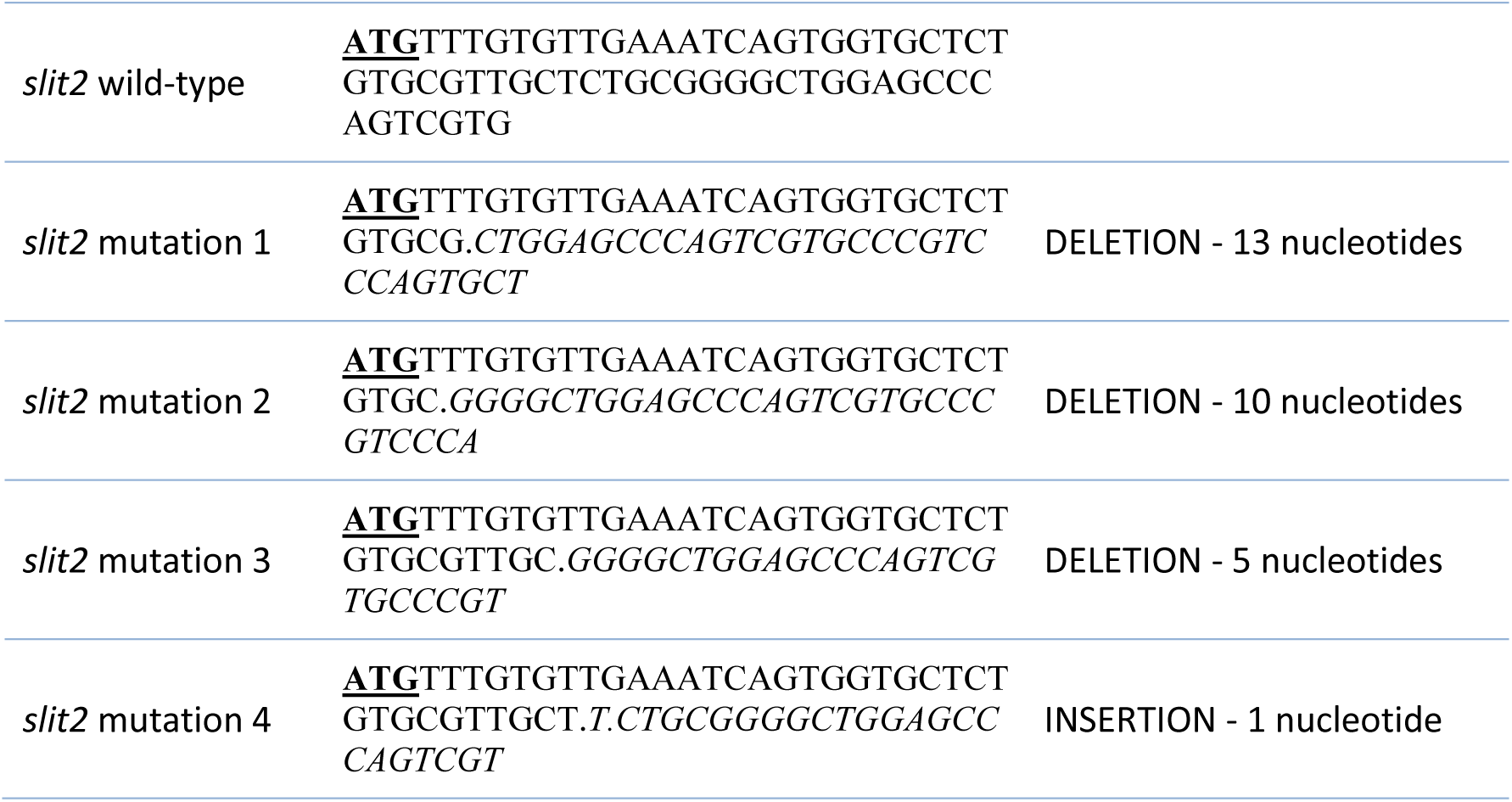

